# N-Glycans Modulate HIV-1 Env Conformational Plasticity

**DOI:** 10.1101/2025.03.26.645577

**Authors:** Mohamed Shehata, Lorenzo Casalino, Madeleine Duquette, Siyu Chen, Alex Flaherty, Elizabeth Villa, Rommie E. Amaro

## Abstract

Human immunodeficiency virus-1 (HIV-1) remains a global health crisis, with over 39 million people living with the virus and no effective vaccine available. Central to HIV infection and immune evasion is the envelope glycoprotein (Env), a heavily glycosylated class I fusion protein that mediates viral entry and is the sole immunogenic target. Despite the recent advancements provided by imaging techniques, the characterization of Env’s structure and dynamics within its native membrane environment remains incomplete. Here, we present microsecond-long, all-atom molecular dynamics simulations of the full-length Env glycoprotein embedded in a biologically relevant lipid bilayer with a complete glycosylation profile. Our simulations reveal a pronounced tilting motion of Env relative to the membrane, with supporting evidence from cryo-electron tomography, which also captures Env tilting within the native membrane. Importantly, we identify a critical role for N-linked glycans at N88 and N611 in modulating the tilting transition. These findings highlight the dual role of Env’s glycan shield as both a protective barrier against neutralizing antibodies and a structural modulator of conformational plasticity. While providing an atomically detailed view of Env in a native membrane environment and advancing the general understanding of its glycan shield and its vulnerabilities, this work also suggests a possible strategy to modulate Env’s conformational plasticity.

## Introduction

Human immunodeficiency virus-1 (HIV-1) remains a global health crisis, having caused between 35.7 and 51.1 million deaths, with over 39 million people currently living with HIV-1 worldwide^1^. The ongoing lack of an effective vaccine continues to hinder global efforts to control the epidemic, which sees approximately 1.5 million new infections annually^1^. At the core of HIV research efforts is the envelope glycoprotein (Env), which mediates viral entry into host cells and serves as the sole viral target exposed on the virion surface for broadly neutralizing antibodies (bNAbs)^2,3^. During viral assembly, Env is expressed as a trimeric gp160, which is subsequently cleaved to become a trimer of gp120-gp41 heterodimers. The gp120 subunit is responsible for binding to the host cell’s CD4 receptor and chemokine coreceptors (CCR5 or CXCR4), while gp41 primes membrane fusion^2,4,5^. Each Env protomer is tethered to the viral envelope via the membrane-proximal external region (MPER), which connects the ectodomain to a single-pass transmembrane domain (TMD) and an intracellular C-terminal domain (CTD)^6,7^. Similar to other class I viral fusion glycoproteins^8^, Env is extensively glycosylated^9–14^, with glycans accounting for more than half of its total mass^12,15,16^. Glycans are essential for Env’s folding, stability, and dynamics^16–19^, while also forming a “glycan shield” that masks underlying proteinaceous epitopes and allows the virus to thwart the host’s immune response^13,16,20,21^. Over the past decade, mass spectrometry-based methods have driven significant progress in the study of the HIV-1 Env glycan shield^9–14^. However, the presence of 23–30 N-linked sites on the gp160 monomer with remarkable microheterogeneity at each site, the glycans’ inherent flexibility, and the vast sequence diversity among circulating viral strains pose significant hurdles for the structural and functional characterization of the glycan shield^10^. These obstacles are compounded by persistent challenges associated with resolving Env’s three-dimensional structure. Low expression levels and poor long-term stability of native Env, coupled with its structural plasticity, have steered most structural research efforts toward the routine use of engineered, truncated versions of the trimer ectodomain, known as SOSIP trimers^22^. This approach has dramatically facilitated the acquisition of high-resolution structures of the Env ectodomain and provided detailed insights into its conformational dynamics^13,23–31^. However, a comprehensive understanding of Env structure in its native viral membrane context has remained elusive^6^.

Obtaining high-resolution structures of the MPER, TMD, and CTD is challenging due to their proximity to the viral membrane^32^. The structures of these regions have only been resolved in isolation from the ectodomain using nuclear magnetic resonance (NMR) ^32–37^. Nonetheless, the majority of current knowledge about Env’s *in situ* structural plasticity has been primarily derived from studies employing cryo-electron tomography (cryo-ET) and cryo-electron microscopy (cryo-EM)^6,7,38^. For example, Prasad et al.^6^ reported Env tilting on the native membrane in an unliganded state using cryo-ET. Similarly, Env tilting was also observed upon interaction with MPER-directed antibodies, as demonstrated via cryo-EM.^7^ In the same study, the authors refer to Env tilting as a part of the Env neutralization mechanism for such antibodies.^7,39^ While these studies shed light on Env’s *in situ* dynamics, their resolution constraints hinder the ability to capture fine structural details. Key aspects, such as the molecular mechanism underlying large conformational transitions, like Env tilting, and the dynamics of the glycan shield, cannot be adequately captured by current imaging techniques. To overcome these limitations, molecular dynamics (MD) simulations provide high-resolution, atomic-level insights, enabling detailed characterization of conformational changes closely tied to key functional processes or immunogenic properties. In this regard, MD simulations have been successfully used to investigate the dynamics of class I fusion glycoproteins, either *in situ* or embedded in a native membrane, for SARS-CoV-2 spike^40–46^ and influenza hemagglutinin^47,48^. On the other hand, computational research addressing Env dynamics remains relatively sparse and underexplored. Some studies have examined the dynamics of the truncated membrane-interacting regions (TMD^49^ or MPER–TMD^37^ without the ectodomain) or a single gp120 monomer with^18^ or without glycans^50^. Other studies have focused on the ectodomain trimer with a simplified glycosylation profile^51^, or investigated an unglycosylated full-length model embedded in a nanodisc membrane through coarse-grained simulations.^38^ This narrow range of studies further underscores the need to bridge the critical gap in understanding the dynamics of the full-length HIV-1 Env in its native membrane environment.

Here, we present µs-long, all-atom, explicitly solvated molecular dynamics (MD) simulations of the full-length Env glycoprotein in the closed state embedded in a native membrane lipid bilayer, with a complete glycosylation profile consistent with glycoanalytic data (Fig. 1). Our simulations reveal that Env undergoes a pronounced tilting motion relative to the membrane normal, exhibiting extraordinary flexibility that is integral to its activity. Consistent with MD simulations, structural characterization of Env expressed in enveloped virus-like particles (eVLPs) using cryo-electron tomography (cryo-ET) further demonstrates Env’s intrinsic dynamic nature, highlighting its ability to adopt multiple orientations relative to the viral membrane.

**Figure 1.**
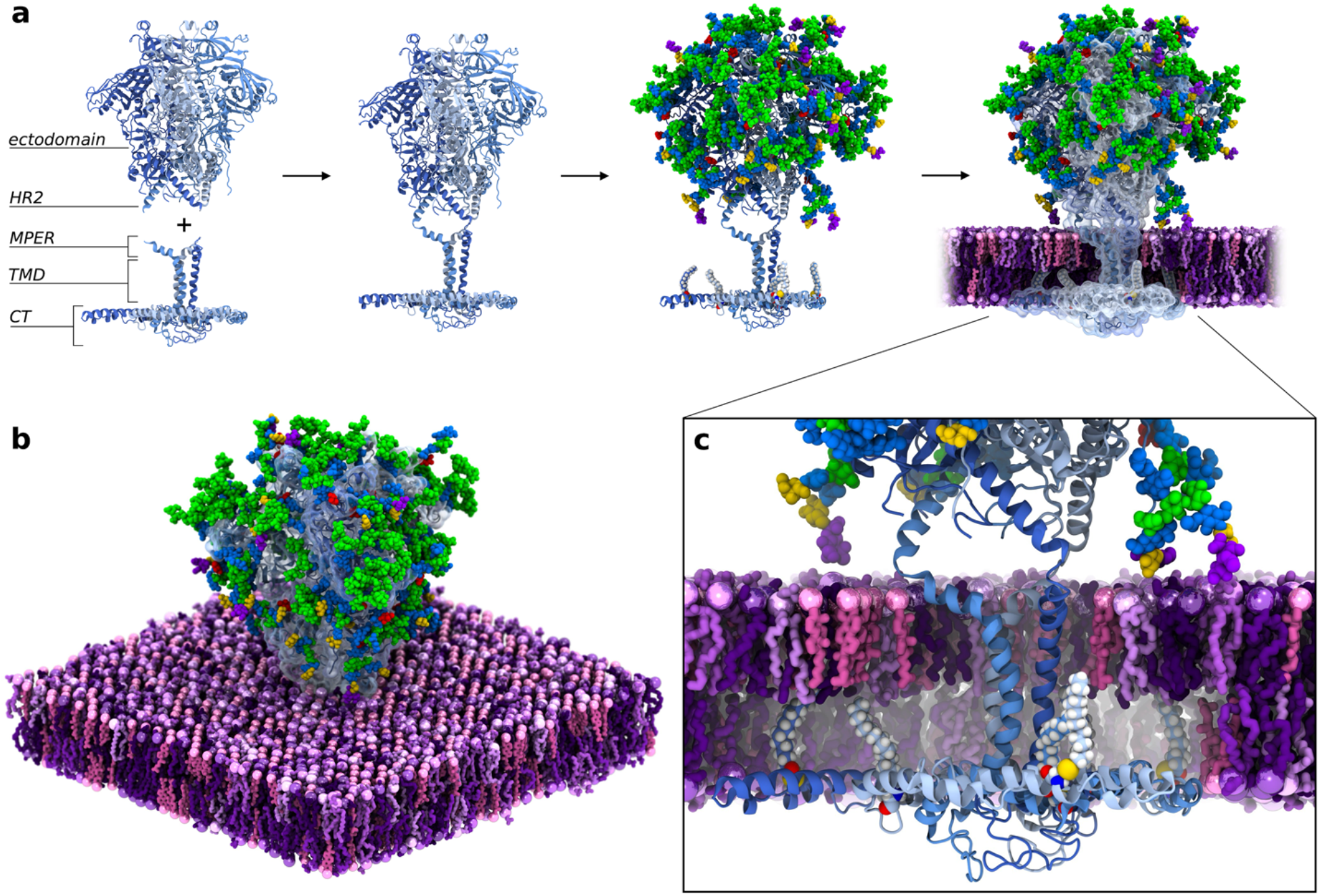
Modeling and simulations of the full-length, fully glycosylated HIV-1 Env trimer in the closed state. (**a**) Workflow of the assembly of the ectodomain, MPER, TMD, and CT domains into the full-length, fully glycosylated model of the HIV-1 Env trimer embedded into an all-atom membrane bilayer mimicking the composition of the viral budding site. The protein is depicted with light blue cartoons, with each protomer differentiated by distinct shades. N-glycans are shown in van der Waals (vdW) representation: GlcNAc in blue, mannose in green, fucose in red, galactose in yellow, and sialic acid in purple. Lipid headgroups are depicted as van der Waals (vdW) spheres, with shades of purple (from darker to lighter) representing DPPC, POPE, POPS, PSM, and POPC, and pink indicating CHOL. Lipid tails are shown in licorice representation. (**b**) Oblique view of the fully assembled, glycosylated Env trimer embedded into an all-atom membrane bilayer (**c**) Close-up view of the Env’s membrane-interacting region, where the palmitoylated cysteines within the CT are depicted with vdW spheres.

Beyond shielding, our simulations suggest that two N-linked glycans, N88 and N611, may play a structural and functional role in modulating the conformational transitions between the Env’s upright and tilted states. Driven by coupled hinging and scissoring motions, MPER and TMD are found to adopt a diverse set of conformations pivotal to Env tilting, some of which show increased accessibility to an MPER-directed antibody. Finally, we provide an atomically detailed view of the Env’s glycan shield over microsecond time scales. The Env’s thick, sugary cloak demonstrates a remarkable ability to restrict access to larger molecules, such as neutralizing antibodies. Overall, the simulations and the tomographic data presented here enhance and expand the existing structural and biological data, offering a previously unexplored, atomic-level perspective on the full-length Env glycoprotein’s structure and dynamics in a native membrane environment. The identified glycans play dual roles as both shielding devices and functional modifications integral to Env’s conformational changes. Our findings could suggest a potential strategy to modulate the conformational plasticity of Env, holding the potential to pave the way for the development of effective vaccines and therapeutic interventions.

## Results

### N-Glycans at N88 and N611 modulate Env tilting cooperatively

In this work, we constructed and simulated a full-length, fully glycosylated model of the HIV-1 Env trimer. The model was built in three different steps, as fully detailed in the Methods section in the Supplementary Information (SI) and illustrated in Fig. 1a, b. The model includes the ectodomain region (residues 631–662), based on the BG505 SOSIP.664 X-ray structure (PDB ID: 5T3Z^23^), and the membrane-interacting region (residues 663–856, comprising MPER, TMD, and CT), modeled from the NMR structure with PDB ID: 7LOI^37^. We note that stabilizing mutations specific to the SOSIP design were reverted to reflect the native sequence of clade A (BG505) Env. Residue numbering referenced throughout this work aligns with the HBX2 sequence, as reported in Fig. S1. The resulting construct was glycosylated at all 69 N-glycosylation sites (23 N-linked sequons per monomer across three monomers) using an asymmetric, site-specific glycoprofile consistent with glycoanalytic data (see Supplementary Tables 1–3 for composition)^9–11^. Following cysteine palmitoylation within the CT (Fig. 1a, c), the resulting model was embedded into an all-atom membrane bilayer mimicking the composition of the viral budding site (see Supplementary Table 4). Explicit water molecules and ions were added, yielding a system of ∼1.3 million atoms.

We performed µs-long MD simulations to investigate the conformational dynamics of the Env glycoprotein, conducting five independent replicates to ensure robustness and reproducibility. Remarkably, our simulations revealed a pronounced tilting motion of the Env ectodomain across multiple replicates (Fig. 2a and Supplementary Movie 1). We quantified the extent of this motion throughout the simulations by calculating the tilt angle of the Env ectodomain relative to the membrane normal (Fig. 2a, b). In four out of five instances, the Env rapidly tilts by ∼30°, occasionally reaching tilt angles of ∼40° (Fig. 2b). In only one case, the Env oscillates around its initial upright conformation, remaining within the 0-10° range.

**Figure 2.**
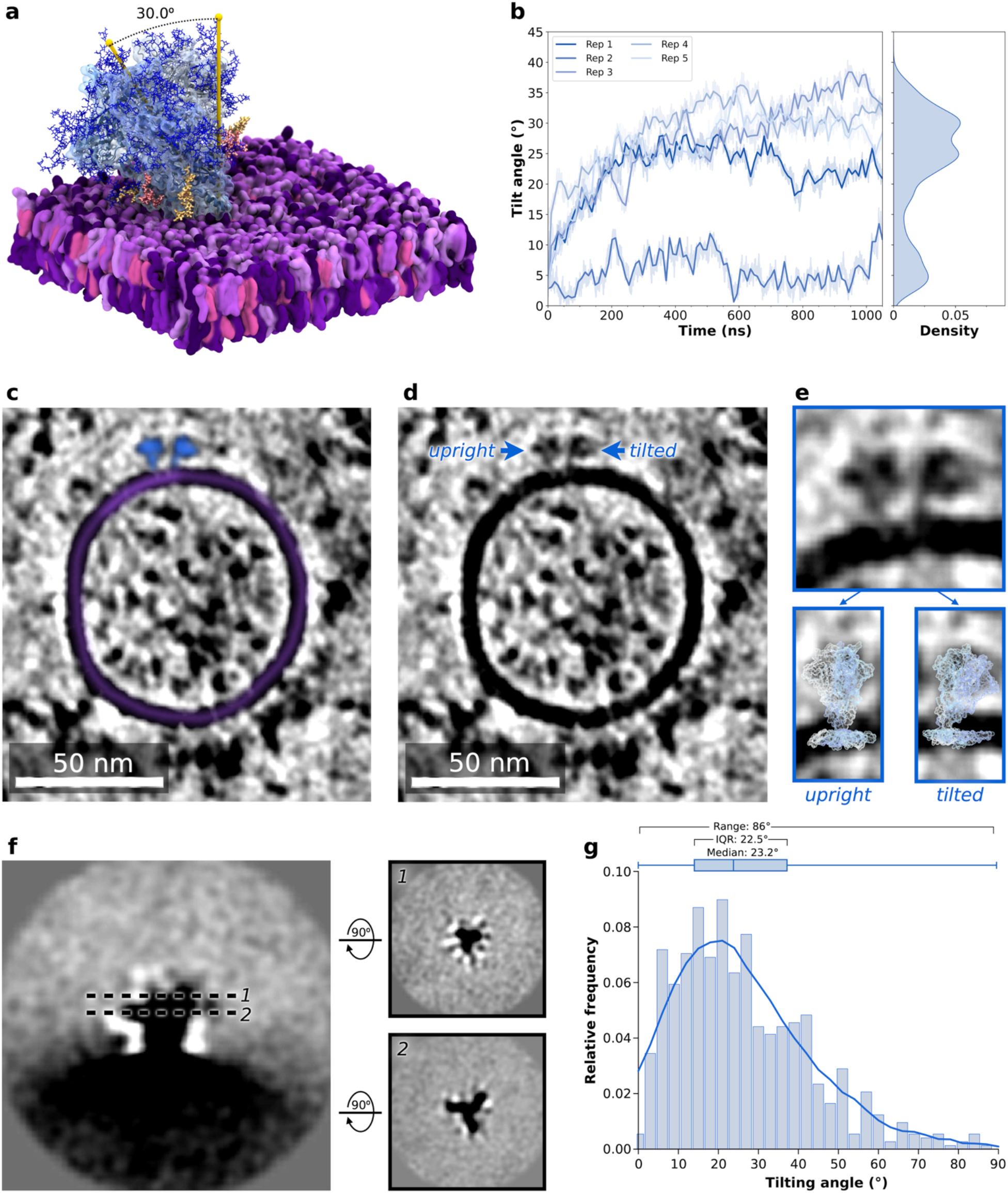
MD simulations reveal pronounced Env tilting motion. (**a**) A representative snapshot of the Env protein showing significant tilting motion relative to the membrane. The Env is displayed with a transparent blue representation, the membrane with a purple surface, and N-glycans are shown with thin blue sticks. N-glycans N88 and N611 are colored in yellow and red, respectively. A tilt angle of ∼30.0° is indicated by drawn yellow cylinders. (**b**) The plot on the left displays the time evolution (x-axis, ns) of the Env tilt angle (y-axis, degree) across the five independent MD simulation replicates. The plot on the right shows the kernel density distribution (x-axis) of Env tilt angles (y-axis, degree). (**c**) Deconvoluted tomogram slice of a virus-like particle showing Env (blue) coating the membrane surface (purple). (**d**) Same tomogram slice as (c) with representative upright and tilted Envs indicated. (**e**) Top: Inlay is a closer look at two Envs. An upright Env is on the left and a tilted Env is on the right. Bottom: Upright and tilted conformations of Env extracted from MD simulations are overlayed on the respective upright and tilted Envs shown in the panel above. (**f**) Different slices (dashed lines labeled as 1 and 2) of the Env average density from the x-y (left) and x-z (right) view. (**g**) Histogram and box-and-whisker plot of the Env tilting distribution from the cryo-ET dataset. Values are binned by rounding to the nearest multiple of 3. Range, interquartile range, and median values are shown.

To experimentally assess Env’s conformational flexibility, we incorporated Env into the envelope of virus-like particles (eVLPs) to preserve its native membrane topology, using the EABR technology described by Hoffmann et al.^52^ to enhance its incorporation efficiency. After purification, the eVLPs were prepared for cryo-ET and 705 tilt series were collected (see Methods in the SI). The reconstructed tomograms were manually inspected, with each Env identified and marked for subtomogram extraction and subsequent analysis (Fig. 2c–f and Fig. S2). Cryo-ET reveals multiple orientations of Env, demonstrating its conformational flexibility *in vitro* (Fig. 2d, e). To calculate the range of motion empirically, the orientation of Env after refinement was used to geometrically determine the tilt angle relative to the normal plane of the eVLP membrane (Fig. S3). The distribution of the Env tilt angles indicates that unbound Env has an interquartile range of 22.5° and a median tilt angle of 23.2° (Fig. 2g), aligning with our simulations, where most tilted conformations fall within the 25°–35° range (Fig. 2b). Overlay of upright and tilted conformations extracted from MD simulations shows remarkable agreement with Env densities identified from tomogram slices (Fig. 2e). We note that the tilt angles determined from cryo-ET subtomogram analysis exhibit a broad distribution peaking around 10–30° (Fig. 2g), whereas our simulations reveal a more defined bimodal distribution, with one peak corresponding to the upright state and a predominant peak for the tilted state (Fig. 2b). This difference likely arises from the inherent limitations of conventional MD simulations, where the accessible timescales restrict the number of observed tilting events. In our simulations, Env typically undergoes a single tilting event that progresses quickly without fully returning to the upright orientation (Fig. 2b). In contrast, experimental measurements sample a broader range of conformations over longer timescales, allowing for multiple independent tilting events (Fig. 2e). Moreover, the absence of the natural membrane curvature in our lipid bilayer model likely results in an underestimation of Env tilting. In simulations with a more realistic *in situ* setting,^48^ the local membrane convexity would likely allow sampling of Env’s tilted conformations exceeding 40°, as captured by cryo-ET (Fig. 2g). Consequently, cryo-ET data obtained here offer a more comprehensive view of Env’s conformational landscape, while the simulations provide atomistic insights into the molecular mechanism of tilting. Notably, our findings are consistent with previous structural studies that reported tilting in both liganded and unliganded forms of Env.^6,7,39^ For instance, Prasad et al.^53^ observed Env tilting in the unliganded state using cryo-ET; however, Env’s full range of motion, a critical aspect that we address here through both simulations and cryo-ET, had not been characterized in that study. Similarly, Richard et al.^39^ investigated Env’s tilting in its complex with CD4, reporting a median tilt angle of 22.7°, which closely resembles what we report here for Env in the unbound state. This similarity suggests that Env tilting is an intrinsic property of Env, occurring independently of receptor binding. By providing a more detailed and quantitative analysis of Env’s mobility, our work bridges these previous observations, offering new insights into the extent and potential functional relevance of Env tilting.

Visual inspection of our simulations revealed direct interactions between Env’s N-linked glycans proximal to the gp120-gp41 interface and the lipid membrane (Fig. 2a and Supplementary Movie 1). To quantify these interactions, the contact frequency of each glycan with the membrane was calculated over the course of the simulations (see Methods in the SI). Two glycans, N88 and N611, exhibit the highest contact frequencies with the membrane, with N88 at 58.1% and N611 at 67.0% (Fig. S4). No notable contact frequencies are observed for the remaining glycans; however, interactions with the membrane increase for some of these glycans toward the end of the simulation as a result of Env tilting (Supplementary Table 6). Given the high contact frequency of N88 and N611 glycans, we investigated their conservation among different HIV strains. Interestingly, we found that these two glycans are strongly conserved (Figure S5). Altogether, these observations suggest that N88 and N611 glycans may play a critical role in modulating Env’s structural plasticity, particularly its tilting ability. This discovery builds on prior evidence highlighting the critical role of N88 and N611 glycans in shaping Env’s antigenicity^54–56^. In detail, previous studies showed that N88 and N611 contribute to the binding of antibodies targeting the gp120-gp41 interface epitope^54–56^, with their removal affecting the neutralization potency of such antibodies. Our findings further emphasize the multifaceted roles of these two glycans as we demonstrate their involvement in modulating Env’s tilting. This affects Env’s antigenicity, as tilting, in turn, can alter the accessibility of the MPER epitope. However, additional assays are warranted to corroborate the specific role of N88 and N611 in regulating Env’s tilting transitions and their broader implications for immune recognition.

To explore how N88 and N611 glycans mediate Env tilting, we sought to dissect the frequency and nature of their interactions with the membrane (Fig. 3a).

**Figure 3.**
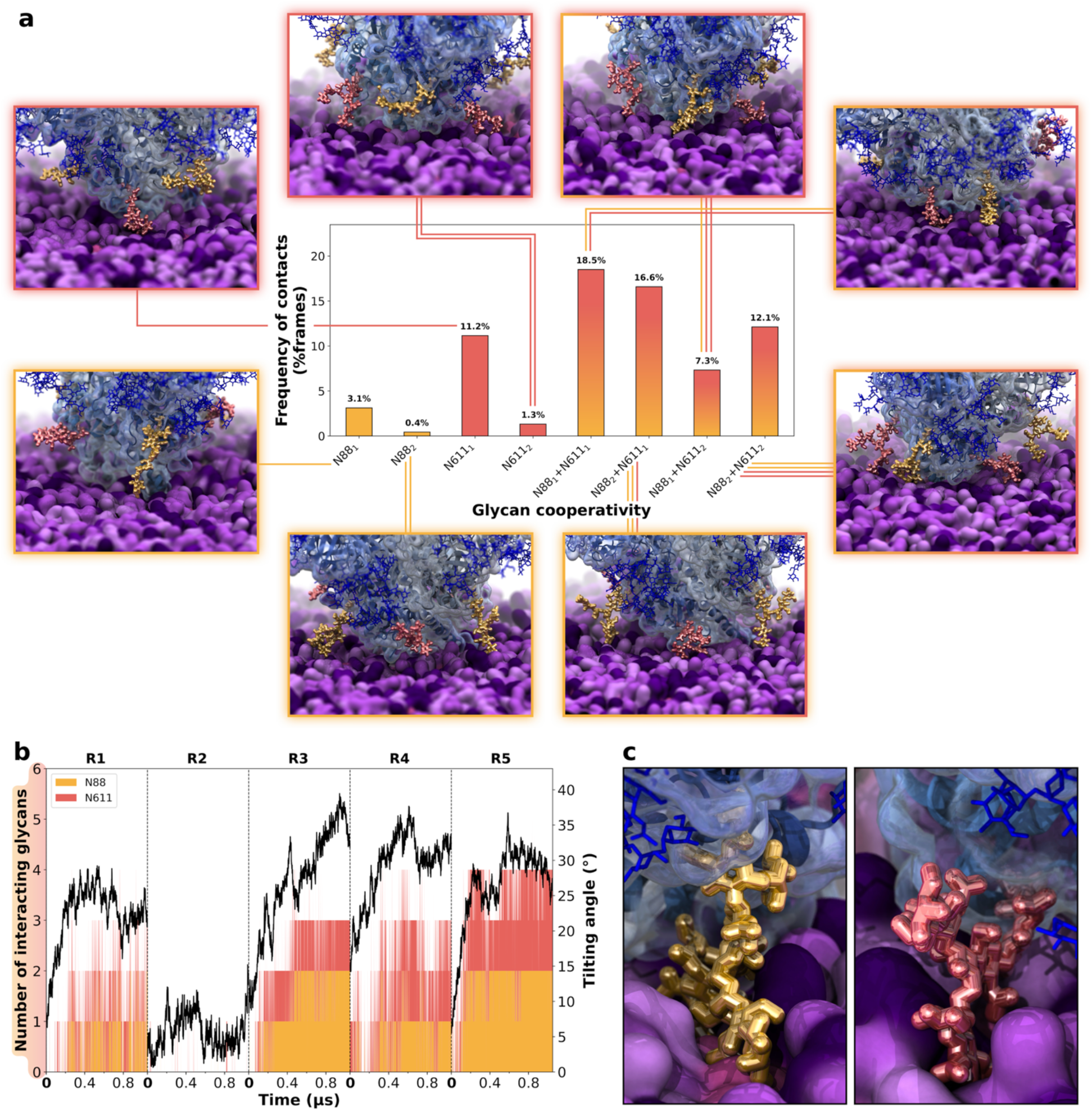
Dissection of N88 and N611 interactions with the membrane. (**a**) The bar plot highlights the frequencies of single (N88_1_ or N611_1_) or combined (N88_2_, N611_2_, N88_1_+N611_1_, N88_2_+N611_1_, N88_1_+N611_2_, N88_2_+N611_2)_ contacts of N88 and N611 glycans with the lipid bilayer accompanied by molecular representations of all interaction combinations surrounding the plot. Contact frequencies were calculated across all simulations and protomers and, for each category, are expressed as % frames with at least one contact from that category. Env is depicted with a blue surface representation, N88 and N611 glycans are shown with yellow and red sticks, respectively, whereas the membrane is displayed with a purple surface. (**b**) Time evolution of the number of N88 (yellow lines) and N611 (red lines) glycans interacting with the membrane (left y-axis), along with the corresponding Env tilt angle in degree (black line, right y-axis) across simulation replicas. (c) Molecular representation of glycans N88 (yellow sticks) and N611 (red sticks) inserted into the lipid bilayer, illustrating their anchoring role during Env tilting.

Our analysis reveals that these glycans contact the head groups of the lipid molecules in diverse ways: often independently, more rarely in combination with the glycan linked to the same site but on an adjacent protomer, and more frequently cooperatively, where N88 and N611 are simultaneously engaged with the membrane. As shown in Fig. 3a, individual interactions exclusively involving only one N88 glycan (N88_1_) or one N611 glycan (N611_1_) exhibit frequencies of 3.1% and 11.2%, respectively, while interactions involving either two N88 glycans or two N611 glycans at the same time are rare (e.g., N88_2_ at 0.4% and N611_2_ at 1.3%). In contrast, for the majority of the simulation time, N88 and N611 concomitantly interact with the membrane. For example, the combination of one glycan from N88 and one glycan from N611 (N88_1_+N611_1_) is observed in 18.5% of the frames, the highest frequency among all combinations. Similarly, other cooperative interactions, such as two glycans from N88 and one glycan from N611 (N88_2_+N611_1_) or one glycan from N88 and two glycans from N611 (N88_1_+N611_2_), display notable frequencies of 16.6% and 7.3%, respectively. To further illustrate these scenarios, we have visualized all observed glycan interaction combinations in Fig. 3. Overall, these observations emphasize how the synergy between N88 and N611 glycans is crucial for modulating Env tilting, with the different interaction patterns actively shaping the tilting orientation and propensity.

Next, to gain further insights into the mechanism of Env tilting, we tracked the aggregate number of N88/N611 interacting glycans across the three protomers throughout the simulations, with a particular focus on the relationship with Env tilting. As shown in Fig. 3b, at least one glycan from N88 or N611 engages with the lipid bilayer at the onset of Env tilting. As tilting progresses, the number of N88/N611 interacting glycans steadily increases, reaching up to four. Notably, this pattern is absent in simulation R2, where no significant tilting occurs. Furthermore, Fig. 3c provides a molecular visualization of N88 and N611 glycans deeply embedded into the lipid bilayer, suggesting a role as conformational anchors during Env tilting. Transient anchoring, where N88 and N611 insert deeply into the lipid outer leaflet, likely stabilizes the tilted conformation of Env by providing further structural support to the points of contact with the membrane. Altogether, these observations suggest that glycan interactions at N88 and N611 are not only critical for initiating Env tilting but also for stabilizing the tilted conformations. Interestingly, this dynamic behavior, mediated by hinge regions and glycan interactions, appears to be a shared feature across class I viral fusion proteins, including SARS-CoV-2 spike^40,41,43,44,46,57^, and influenza hemagglutinin^48,58^. These proteins leverage conformational flexibility and glycan-mediated stabilization^40,46,59^ to optimize host cell engagement, fusion, and immune evasion. The presence of these conserved mechanisms underscores their functional importance in viral infection while also highlighting potential conformational targets for the development of therapeutics against them.

### Tilting of Env trimer is accommodated by MPER and TMD hinging motions

The MPER (residue 660 to 683) and the TMD (residue 684 to 706) are critical structural domains of Env, playing critical roles in membrane anchoring, fusion, viral entry, and structural stabilization of the Env trimer^7,60^. However, as detailed in the introduction, their dynamics within the full-length Env and native viral membrane context remain largely underexplored, leaving critical gaps in understanding their contributions to the full spectrum of Env structural dynamics, including tilting dynamics. The MPER bears a hinge-like structure, with a linker loop serving as the pivot point between an N-terminal α-helix (spanning the last three helical turns of HR2) and a three-turn C-terminal α-helix positioned at the level of the membrane’s phosphate head groups (Fig. 4a)^61^. This three-turn helix transitions into the single-pass transmembrane helix, i.e., the TMD, forming an almost continuous helix slightly bent at residue 683 (Fig. 4a). Consequently, this structural organization defines two hinge points, which are illustrated in Fig. 4a: one at the linker loop in the middle of the MPER, hereafter referred to as ‘MPER hinge angle,’ and another at residue 683, marking the transition between the MPER and TMD, hereafter referred to as ‘MPER– TMD hinge angle.’

**Figure 4.**
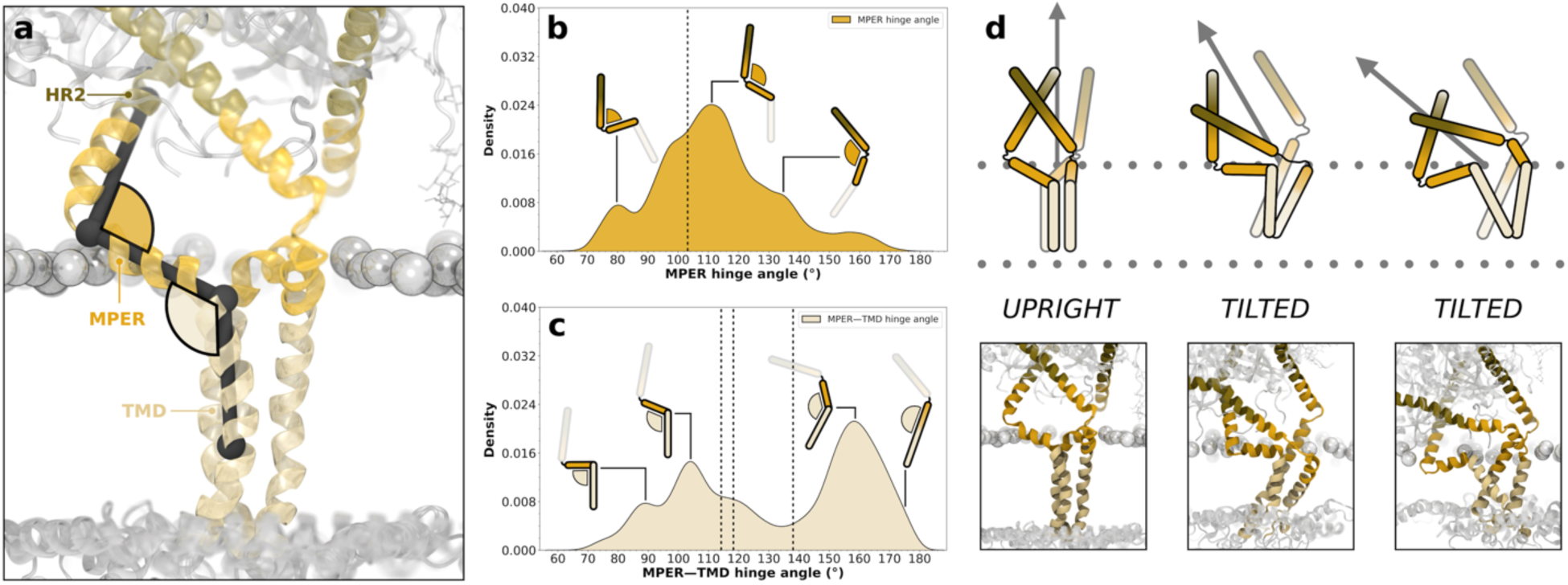
Conformational landscape explored by MPER and TMD. (**a**) Schematic representation of the two hinge angles analyzed: MPER hinge angle (orange) and MPER–TMD (beige) hinge angle. (**b**) Kernel density distribution (y-axis) of the MPER hinge angle (x-axis, degree). (**c**) Kernel density distribution (y-axis) of the MPER–TMD hinge angle (x-axis, degree). The dashed lines indicate the initial angle values derived from the starting model before simulations. (**d**) Representative conformations from simulations and relative schematic illustrations showing the upright and tilted states of the Env trimer, highlighting MPER and TMD adaptations as the trimer transitions to tilted orientations relative to the membrane.

Using kernel density estimates, we analyzed the distributions of these hinge angles across our simulation replicates to capture the range of conformations sampled (Fig. 4b, c). The distribution of MPER hinge angles shows a broad peak centered around 110°, spanning a range of angle values from 75° to 165° (Fig. 4b). Schematic representations depicted above the distribution plot in Fig. 4b illustrate the most significant conformations sampled by the MPER. At smaller angles (∼85°), the MPER adopts a more tightly bent state. Around the peak angle (∼110°), it maintains a slightly open yet stable configuration, consistent with the initial conformation (indicated by the dashed line). This suggests that the MPER retains its starting hinge-like structure while exhibiting flexibility needed for its functional roles, such as mediating membrane interactions and supporting tilting transitions. At larger angles (∼140°), the MPER transitions into a more extended configuration, stretching along the linker loop. This analysis may indicate that there might be structural or energetic constraints limiting extreme bending or extension.

Differently from the MPER hinge angle, the MPER–TMD hinge angle distribution exhibits a bimodal profile, with two major peaks centered at approximately 105° and 160°, reflecting the predominant conformational states sampled during the simulations (Fig. 4c). The schematics above the plot illustrate the key sampled conformations corresponding to specific regions of the distribution. At ∼90°, the MPER–TMD hinge is in a tightly bent, compact configuration. At ∼105°, the MPER–TMD helix transitions to a curved, moderately bent conformation. At ∼160°, the hinge assumes an extended configuration, with a slight bend in correspondence of residue 683, whereas at ∼180° it reaches a super extended conformation where the bend is no longer present (Fig. 4c). The dashed line in the plot in Fig. 4c represents the values of the MPER–TMD hinge angle as in the initial conformation of our model (Fig. 4a), which, in turn, is based on NMR data lacking the ectodomain and membrane^37^. The sampled conformations observed during our simulations diverge significantly from this initial conformation, highlighting the limitations of NMR-based models in accurately capturing TMD dynamics under biologically relevant conditions. Without the ectodomain and membrane, the NMR-based model is unable to account for the broader conformational space accessed by the MPER–TMD hinge. In contrast, our simulation captures a more comprehensive range of hinge dynamics, including both bent and extended conformations, demonstrating the advantages of integrating environmental and structural components into the model. This comparison underscores the importance of simulations in resolving the full conformational landscape and overcoming the limitations inherent in NMR-based structural approaches.

To summarize the conformational changes occurring at the level of the MPER and TMD regions during Env tilting, we integrated the information derived from the distribution of the two hinge angles with representative structural snapshots extracted from the simulations (Fig. 4d). This approach enabled us to outline the mechanism by which the MPER and TMD accommodate or facilitate Env tilting. Their combined conformational transitions between compact, bent, and extended configurations are affected by the extent of tilting and the trimeric spatial arrangement. The schematic illustrations reported in Fig. 4d, along with the corresponding simulation snapshots below, summarize some of the conformational changes taking place at the level of the MPER and TMD within the trimeric Env structure, illustrating both the upright and tilted states. In the upright state, represented in this case by the initial conformation of our model, all three protomers exhibit symmetric hinge angles. The MPER hinge angles are ∼110° (as indicated by the dashed line in Fig. 4b), while the MPER–TMD hinge angles correspond to ∼110°/∼135° (as indicated by the dashed line in Fig. 4c). This state represents the starting configuration of the trimer prior to tilting. In the tilted state, asymmetry emerges among the three chains as the trimer adjusts its hinge angles. The MPER of the protomer located in the direction of the tilt bends further, reducing its hinge angle to ∼95° or even ∼80°, corresponding to the conformation highlighted by the left shoulder peaks in the distribution in Fig. 4b. Concurrently, the MPER–TMD hinge adopts a more bent angle (∼90° or ∼100°), aligning with the two peaks on the left-hand side of the distribution in Figure 4c. In contrast, the other two protomers extend to accommodate the tilt, with their MPER hinge angles ranging from ∼135° to ∼160, as indicated by the right shoulder peaks in the distribution reported in Fig. 4b. Similarly, their MPER–TMD hinge angles extend to ∼160° or ∼180°, aligning with the two peaks on the right-hand side of the distribution in Fig. 4c. In agreement with previous hypotheses suggesting a loosely folded scissoring motion of the TMD helices in their ground state^7^, our simulations reveal a significant mobility of the TMD helices within the membrane. Analysis of the angle of intersection between the TMD helices and the membrane leaflets shows that the TMD can shift from a vertical orientation (∼90°) to a tilted orientation (∼40°) (Fig. S6). Although we do not directly quantify the scissoring motion, representative snapshots from our simulations demonstrate that the TMD helices orient themselves in a tilted fashion within the lipid bilayer (Fig. 4d), forming an X-like arrangement with adjacent protomers. We hypothesize that this dynamic reorientation of the TMD helices within the bilayer also plays a key role in accommodating Env tilting.

### Atomic-level overview and analysis of Env’s glycan shield

Understanding the strategies that viruses employ to evade the host immune response is crucial for designing effective vaccines^40^. Among these, the glycan shield stands out as one of the most effective mechanisms utilized by class I viral fusion glycoproteins, including HIV-1 Env, to thwart the humoral response^16,20,62,63^. Approximately half of the molecular weight of Env is derived from N-glycans, which are strategically distributed across the ectodomain^8,13,64,65^. These glycans form a dynamic barrier, hampering antibody binding to antigenic regions while selectively allowing access to receptor-binding surfaces necessary for infection^20^. Interestingly, many bNAbs require glycan recognition for high-affinity binding, underscoring the critical role of glycans in modulating Env’s antigenicity^62,63^ and informing immunogen design^66^.

Aiming to characterize the glycan shield of Env, we analyzed its spatial distribution and dynamic behavior during our MD simulations. In Fig. 5a, b, we provide previously unappreciated atomic-level views of the glycan shield, revealing its configuration and appearance over the microsecond time scale. Due to their inherent flexibility and multi-antennary, branched structures, N-glycans are poised to rapidly cover a substantial portion of the underlying proteinaceous residues, acting on timescales much shorter than those required for protein conformational changes. We overlaid equally interspersed glycan conformations sampled over a 1-μs-long simulation onto a reference Env conformation, probing the protein for potential sites of vulnerability. The atomic-level representation of the glycan shield reveals that the Env is almost entirely covered, leaving minimal exposure to proteinaceous residues (Fig. 5a, b). We note that our MD simulations account only for the Env in the closed, unliganded ground state without exploring opening motions leading to the open-occluded state^29^. This dynamic view of the glycan shield further emphasizes the remarkable density of the Env’s glycan shield, especially when compared to the one in the SARS-CoV-2 spike protein^40^.

**Figure 5.**
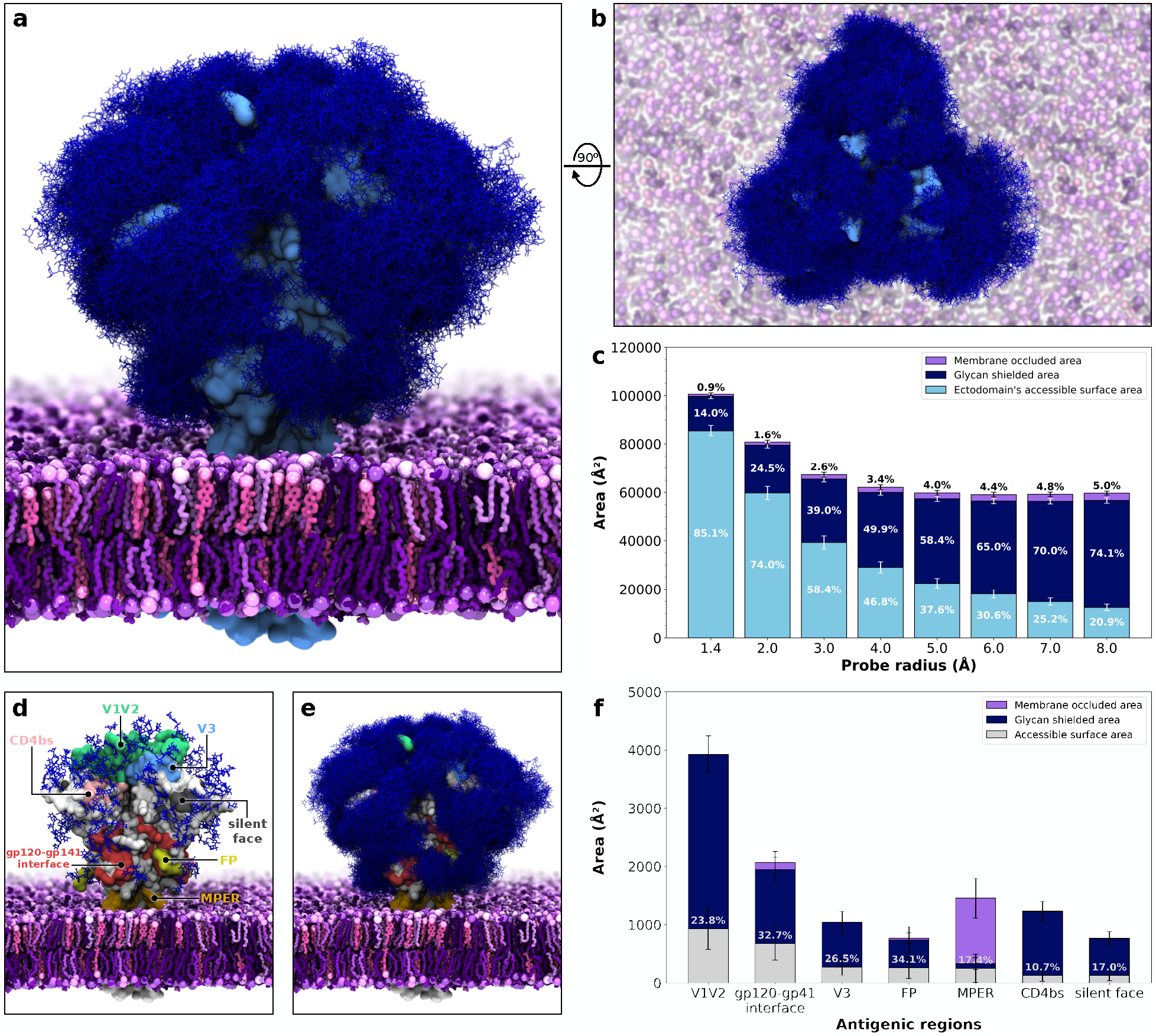
Glycan shield of HIV-1 Env. (**a, b**) Molecular representations of the HIV-1 Env trimer’s glycan shield, shown from a side view (**a**) and a top-down view (**b**). Glycans are shown as a blue mesh ensemble of overlaid conformations sampled during a 1-µs-long simulation. The protein is represented by a cyan surface, while the membrane is depicted in purple, with lipid headgroups represented as van der Waals spheres. (**c**) Quantification of the accessible surface area (cyan), glycan-shielded area (dark blue), and membrane-occluded area (purple) in Å^2^ across probe radii ranging from 1.4 Å to 8 Å (x-axis), with relative percentages indicated within each stacked bar. For each stacked bar, the values have been averaged across replicas, and the error bars indicate the standard deviation of the respective mean value. (**d**) Overview of the HIV-1 Env trimer’s major antigenic regions. The protein is represented with a light gray surface, and the antigenic regions are highlighted in distinct colors: CD4bs (pink), V1V2 (green), V3 (cyan), silent face (dark gray), gp120-gp41 interface (red), fusion peptide (FP, yellow), and the MPER (orange). Glycans are depicted as blue sticks. (**e**) Alternate representation of the Env trimer as in panel D, but with glycans shown as a blue mesh ensemble of overlaid conformations sampled during a 1-µs-long simulation. (**f**) Quantification of the accessible surface area (light gray), glycan-shielded area (dark blue), and membrane-occluded area (purple) for the antigenic regions represented in panels E and F using a probe radius of 7.2 Å. For each stacked bar, the values have been averaged across replicas, and the error bars indicate the standard deviation of the respective mean value.

To assess the glycan shield’s effectiveness in occluding the underlying proteinaceous residues, we calculated Env’s accessible surface area (ASA) to spherical probes with radii ranging from 1.4 Å to 8 Å (Figure 5c, see Methods section in the SI for a complete description). A probe with a radius of 1.4 Å is commonly used to describe a water molecule, larger radii (2.0–5.0 Å) are used to represent small molecules, while 7.0–8.0 Å approximate the size of the complementarity-determining regions (CDRs) of an antibody’s variable fragment (Fv)^67^. This analysis demonstrates a striking dependency of ectodomain accessibility on probe size. At the smallest probe radius of 1.4 Å, approximately 85% of the ectodomain remains accessible, reflecting the potential facility of small molecules to penetrate the glycan layer. As the probe radius increases, the extent of glycan shielding becomes gradually more pronounced. For probes with a 3.0 Å radius, accessibility decreases to ∼58%. At radii (7–8 Å), which approximate the size of an antibody’s CDR, glycans obscure ∼70–74% of the ectodomain surface, leaving only ∼20–25% accessible (Fig. 5c). These findings highlight the general ability of the glycan shield to function as a dynamic barrier predominantly restricting access to larger molecules, such as antibodies, whose binding to epitopes occurs on significantly slower timescales (µs to ms) than glycan conformational fluctuations (ps to ns)^40^. Nonetheless, it is important to note that the glycan shield must adapt to allow access to the ectodomain surface during the infection process^68^. Our molecular representation of the glycan shield in Fig. 5a,b shows the presence of multiple vulnerabilities, akin to cracks in the sugary armor. Several antigenic regions on Env have been identified as targets of bnAbs, with epitopes primarily located in the variable regions 1 and 2 (V1/V2)^69–72^, V3^69,73,74^, the CD4 binding site (CD4bs)^75–77^, the silent face^78,79^, the gp120-gp41 interface^25,54,80^, the fusion peptide (FP)^54,81^, and the MPER^61,82–84^ (see Fig. 5d, e and Supplementary Table 7 for a detailed list of bnAbs targeting these regions). To unravel the vulnerabilities of these key antigenic sites, we measured the extent of occlusion conferred by both the glycan shield and the membrane (Fig. 5f). This analysis provides a comparative view of how structural shielding varies across epitopes, offering insights into potential vulnerabilities that could be leveraged for vaccine design. For this purpose, we only calculated the accessibility to a probe mimicking the size of an antibody’s CDR (radius 7.2 Å). Our findings indicate that the FP is the most accessible antigenic site, with ∼34.1% of its surface exposed, followed by the gp120-gp41 interface (32.7%). Despite the participation of glycans N88 and N611 in the neutralization mechanism required by bnAbs 35O22^80^ and PGT151^54^, these regions stand out as the most vulnerable to immune recognition. Conversely, the CD4bs is the least accessible, with only 10.7% of its surface exposed, making it the most glycan-shielded antigenic region. Notably, the extensive shielding is due to glycans lining the binding site, mostly N276, even though the site itself is not glycosylated. Our findings align with previous reports indicating that the CD4bs is primarily occluded by the glycan shield in the prefusion-closed state, reinforcing the notion that its limited immunogenicity is due to extensive glycan shielding^85^. V1V2 and V3 exhibit high levels of glycan shielding, with only 23.8% and 26.5% accessibility, respectively (Fig. 5f). This is consistent with the neutralization mechanism of bnAbs targeting V1V2, such as PG9^71^ and PGT145^69^, which strongly depends on interactions with glycans N156 and N160. Likewise, bnAbs such as PGT121^69^ and BG18^73^ require interactions with glycans N137 and N332 to engage the V3 glycan supersite epitope. Notably, glycan N332 is absent in the HIV-1 strain modeled here, suggesting that the nominal accessibility of V3 may be even lower in other strains where this glycan is present. Interestingly, the silent face exhibits the second-lowest level of accessibility (∼17.0%), indicating that it is largely shielded. This is expected as this epitope requires interactions with glycans N262 and N295 for effective antibody binding, as observed with bnAbs like VRC-PG0562^78^. However, given its partial, albeit minimal, exposure, the silent face may represent an underexplored immunogenic region that could become more accessible under specific conformational states or variations in the Env’s glycan shield.

A noticeable point of vulnerability is observed in the vicinity of the membrane-proximal region (Fig. 5c, e). Given the globular shape of the Env trimer and the lack of an extended stalk tethering the TMD to the ectodomain’s head, we hypothesized that the membrane itself might contribute to obstructing access to the membrane-proximal region of the ectodomain. Across all probe sizes, the viral membrane accounted for only ∼0.9–5% of the total occluded ectodomain’s surface area on average (Fig. 5c). While this contribution is minimal when considering the entire ectodomain, it nonetheless bears a more significant impact for regions proximal to the membrane, such as the MPER, where it accounts for over 80% of the total occlusion for a probe of 7.2 Å radius (Fig. 5f). This suggests that, unlike other antigenic regions primarily concealed by the glycan shield, the MPER remains largely occluded due to its proximity to the lipid bilayer, highlighting the presence of structural constraints governing its accessibility and immunogenicity.

### Env tilting modulates MPER peptide epitope accessibility

The MPER is a highly conserved domain essential for viral membrane fusion and a major target for broadly neutralizing antibodies^61,86^. However, its location near the viral membrane (Fig. 6a), flexibility, and partial coverage by the glycan shield present significant challenges for its effective targeting in vaccine and therapeutic development^61,86–88^.

**Figure 6.**
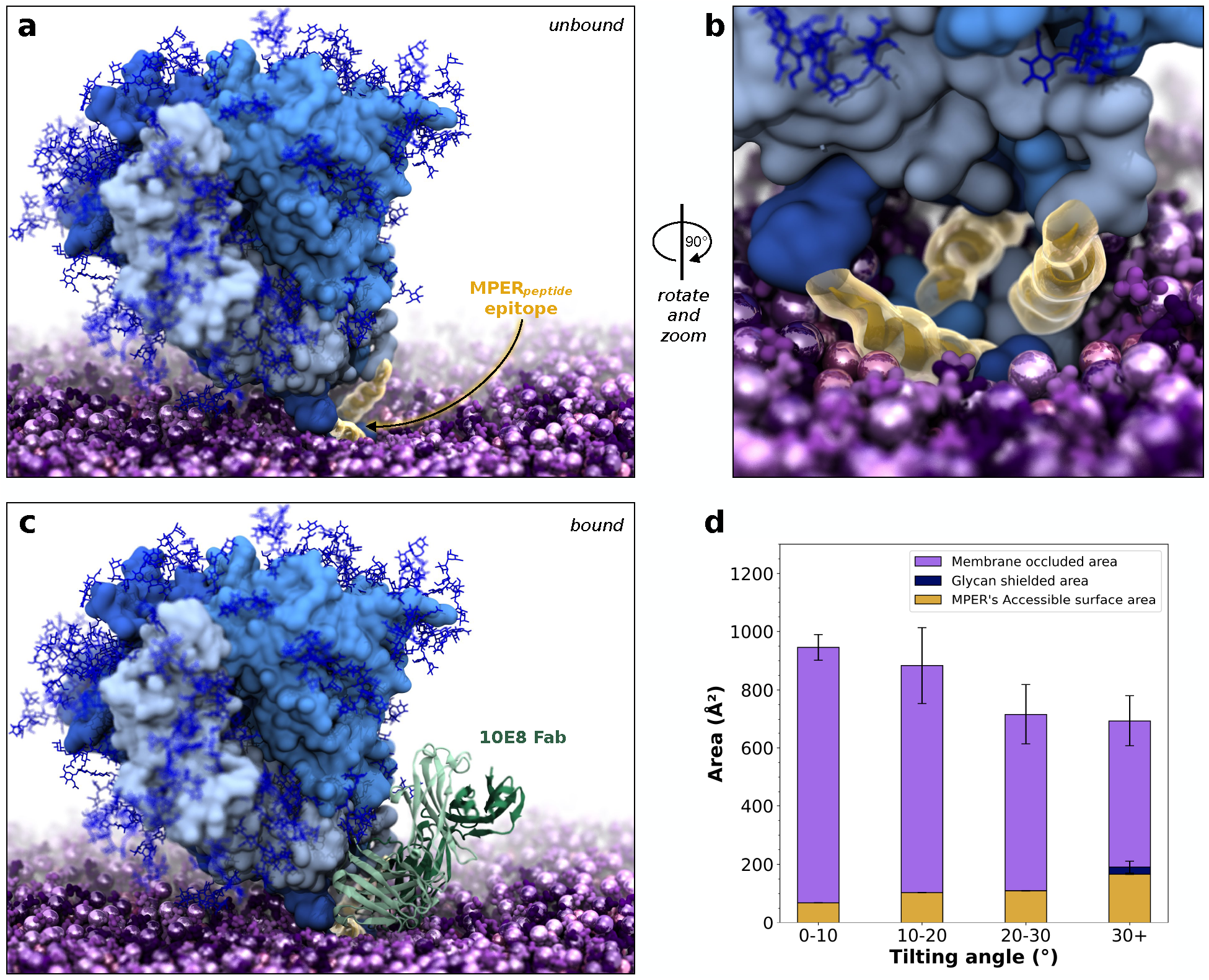
Accessibility of the MPER peptide epitope relative to Env tilt angle. (**a**) Molecular representation of the HIV-1 Env in a tilted conformation shown from a side view, with the MPER peptide epitope (residues 671–683) highlighted in yellow near the viral membrane. N-glycans are depicted as thin, dark blue sticks, while the protein is represented with a light blue surface, with each protomer differentiated by distinct shades. The membrane is depicted in purple, with lipid headgroups represented as van der Waals spheres. (**b**) Rotated and zoomed-in view of the MPER peptide region (yellow), focusing on its spatial arrangement within the viral membrane (purple), with two MPER peptides positioned closer to the viewpoint and prominently exposed, and the third further away buried in the membrane. (**c**) The 10E8 Fab (green cartoons) was docked onto the Env by fitting the MPER peptide from the 10E8 Fab–MPER complex (PDB ID: 4G6F^61^) to the most exposed MPER region sampled in our simulations, demonstrating how it fits into the structural pocket formed by tilting-induced rearrangements of the MPER. (**d**) Quantification of the accessible MPER surface area (yellow), glycan-shielded area (dark blue), and membrane-occluded area (purple) as a function of Env tilt angle (x-axis: 0–10°, 10–20°, 20–30°, 30+°). Relative percentages are indicated within each stacked bar. Error bars represent the standard error of the mean for each area, calculated via bootstrap resampling with 2000 iterations.

The MPER presents several epitopes distributed along its sequence (residues 660–683), capable of eliciting bnAbs with varying levels of potency. Among these, 2F5 predominantly targets the MPER’s N-terminal α-helix (residues 656–671)^84^, Z13e1 mostly binds the MPER’s hinge point (residues 666–677)^83^, 4E10^82^ and 10E8^61^ primarily recognize the MPER’s C-terminal α-helix (671–683), here referred to as MPER peptide epitope. One of the broadest and most potent is 10E8^61^, which was found to be particularly sensitive to residue 681– 683. In Fig. 6a, the Env trimer is shown in a tilted state, illustrating how this conformational change affects the spatial positioning of the MPER. As Env tilts, the MPER peptide becomes increasingly accessible (Fig. 6a, b), underscoring how tilting-induced motions dynamically regulate its exposure. The MPER peptide in the direction of tilting is buried deeper into the viral membrane, reducing its exposure (Fig. 6b). Conversely, the MPER peptides on the opposite side of the tilt are slightly lifted from the membrane, becoming largely unobstructed and increasing their accessibility. The tilt-related conformational change in the MPER forms a pocket-like structural niche, presenting an opportunity for MPER-directed antibodies to engage with the MPER peptide at a more favorable angle of approach. To test whether this pocket could accommodate antibodies targeting the MPER, we aligned the MPER peptide from the 10E8 Fab–MPER complex (PDB ID: 4G6F^61^) to the corresponding, most exposed MPER region sampled in our simulations (visualization of the alignment is reported in Fig. S7). As shown in Figure 6c, the pocket formed by the tilted Env trimer accommodates the 10E8 Fab, allowing it to bind the MPER peptide epitope without any clashes with the membrane or the rest of the Env glycoprotein. This alignment demonstrates how tilting-induced conformational changes in Env can create structural opportunities for antibody binding, even in regions otherwise heavily occluded by the membrane.

To quantify the extent to which Env tilting influences MPER peptide exposure, we clustered all the frames from our MD simulation trajectories into four distinct ensembles based on the degree of Env tilting (0–10°, 10–20°, 20–30°, and 30+°) and calculated the ASA of the MPER peptide epitopes for each ensemble (refer to the Methods section in the SI for a complete description). A probe of a 7.2 Å radius was used to approximate the size of the Fab’s CDR. The obtained values were filtered to include only the most exposed MPER peptide epitope among the three and plotted in Fig. 6d. At lower tilt angles (0–10°), the MPER peptide remains predominantly occluded by the viral membrane, with minimal accessibility for potential interactions. As the tilt angle increases, the extent of membrane occlusion decreases significantly, leading to greater MPER exposure at tilt angles above 30° (Fig. 6d). It is worth mentioning that glycan shielding contributed minimally to MPER occlusion, with the viral membrane playing the primary role in modulating accessibility. These findings suggest potential strategies for vaccine design. The relationship between ectodomain tilting and the accessibility of membrane-proximal epitopes, such as the MPER epitope, underscores the possibility of exploiting tilting-induced conformational changes to enhance the immunogenicity of poorly accessible epitopes. By stabilizing Env in specific tilted conformations or mimicking these dynamics in immunogens, it may be possible to increase the exposure of vulnerable regions, such as the MPER, for antibody recognition. For example, aptly engineered glycans at position N88 or N611 could improve the modulation of tilting dynamics. This approach could be extended to other class I fusion glycoproteins, like influenza HA, where similar mechanisms of membrane-proximal epitope exposure have been observed^48^. Importantly, our simulations reveal that Env tilting can occur independently of, and prior to, antibody binding to the MPER. Previous hypotheses suggested that the approaching antibody must create a wedge between the ectodomain and membrane surface, thereby inducing Env tilting^7^. While we cannot rule out the possibility that antibody binding can also occur in the upright conformation, our findings indicate that Env tilting transiently increases the exposure of MPER epitopes, creating more favorable conditions for antibody binding.

Finally, following the same approach used for the MPER epitope, we probed whether a similar relationship between Env tilting and epitope accessibility was also in place for the gp41-gp120 interface and the CD4 binding site epitopes (Fig. S8). The analysis revealed that, for both epitopes, Env tilting imparts no significant effect on their ASA. In these cases, in contrast to the MPER peptide epitope, the glycan shield remains the dominant contributor to their occlusion. Altogether, these findings emphasize the role of Env plasticity in creating opportunities for antibody binding to otherwise occluded regions, providing valuable insights for the rational design of vaccines and therapeutics targeting the MPER.

## Discussion

In this study, we investigated the conformational dynamics of the HIV-1 Env trimer within its native membrane environment using a combination of all-atom MD simulations and cryo-ET. Our findings reveal that Env undergoes a pronounced tilting motion relative to the membrane normal. Cryo-ET provides an empirical assessment of Env’s full range of motion, revealing a median tilt angle of 23.2°, closely aligning with the tilt range observed in our simulations. Additionally, the similarity between tilt angle distribution observed for the unliganded Env in this study and that reported for Env bound to CD4^39^ suggests that tilting is not induced by receptor binding but rather represents a fundamental aspect of Env’s conformational landscape. Our simulations reveal that Env’s tilting motion is modulated by two N-linked glycans, N88 and N611, upon extensive interaction with the viral membrane. These findings highlight how glycans extend their function beyond shielding to actively regulate Env’s conformational landscape. Importantly, tilting transitions are accommodated by coupled hinging and scissoring motions of the MPER and the TMD, profoundly affecting the exposure of critical immunogenic regions, such as the MPER peptide epitope. Unlike other epitopes where the glycan shield serves as a nearly impenetrable barrier, our investigation reveals that the MPER is primarily occluded by the membrane. Env tilting creates favorable conditions for the binding of MPER-directed antibodies by inducing the formation of an exposed structural niche defined by the MPER positioned on the opposite side of tilting. Beyond providing fundamental insights, these findings might hold significant implications for vaccine and therapeutic development. Aptly engineered constructs designed to modulate tilting dynamics could increase the immunogenicity of the MPER peptide epitope by promoting its exposure, providing a novel strategy for targeting this critical region. Conversely, stabilized Env constructs with reduced tilting might be particularly promising for nanoparticle-based vaccine platforms, where precise epitope positioning can enhance immunogenicity. Furthermore, less dynamic Env constructs could improve high-resolution imaging techniques, such as cryo-EM and cryo-ET, by reducing conformational variability. Such constructs could also reveal druggable pockets, paving the way for novel small-molecule inhibitors. This work exemplifies the power of integrating MD simulations with structural biology techniques, offering actionable insights to inform the next generation of HIV-1 vaccines and therapeutics.

## Methods

An extended description of the modeling procedures, simulation protocols and parameters, analysis workflows and parameters, cloning and expression methods, and cryo-ET data acquisition and analysis is provided in the Supplementary Information. A summary is provided below.

### Methods summary

A full-length, fully glycosylated model of the HIV-1 Env trimer was built by integrating cryo-EM and NMR structural data with loop reconstruction, homology modeling, and AlphaFold2^89^ refinement. The ectodomain was based on the crystal structure of the HIV-1 Env trimer BG505 SOSIP.664 (PDB ID: 5T3Z^23^). SOSIP-specific modifications were reverted to restore the native clade A (BG505) sequence. The membrane-interacting regions (MPER, TMD, and CT) were homology-modeled in i-TASSER^90^ to match the clade A sequence, starting from a clade D NMR structure (PDB ID: 7LOI^37^) as a template. Following HR2 refinement to enable proper alignment and assembly between the ectodomain and membrane-interacting regions, the model was site-specifically glycosylated based on glycoanalytic data^9–11^ and palmitoylated using CHARMM-GUI^91^, then embedded in an asymmetric lipid bilayer mimicking the composition of HIV-1 membrane^92–95^. All-atom MD simulations were conducted using NAMD3^96^ with the CHARMM36m force field^97–99^ in explicit water, with Na⁺ and Cl⁻ ions added to a final concentration of 150 mM. Each system underwent multi-stage equilibration, followed by five independent MD production runs of 1.05 μs each. Analyses included calculation of Env tilt angle, glycan-membrane contacts, hinge and inclination angles, and ASA of the whole protein and specific epitopes.

The full-length BG505 Env gene was cloned into a pcDNA3.1(+) vector, incorporating additional motifs to enhance ESCRT recruitment and promote efficient self-assembly and budding of eVLPs. These eVLPs were expressed in 293FT cells, and purified via ultracentrifugation over a 20% sucrose cushion, and resuspended in 1xPBS before immediate vitrification on cryo-EM grids. Dose-symmetric tilt series of eVLPs were acquired using a Titan Krios G2 microscope. The tilt series were aligned and reconstructed into tomograms, from which Env particles were manually picked for subtomogram averaging in Relion3^100^. Orientations of individual Env particles relative to the VLP surface were then used to calculate tilt angles on a per-particle basis.

## Notes

The authors declare no competing financial interest.

## Data availability

MD simulation data of HIV-1 Env generated in this study (trajectories, initial and final coordinates, input files) will be included in the final version of the article as Supplementary Data and will be made available for download on the Amaro Lab website (https://amarolab.ucsd.edu/data.php#hiv). Cryo-ET map will be deposited in the Electron Microscopy Data Bank (EMDB) under accession code EMD-xxxxx. Raw movies will be uploaded to the Electron Microscopy Public Image Archive (EMPIAR). Source data will be provided in the final version of the article.

## Code availability

This study utilized the standard build of the simulation software NAMD3 (https://www.ks.uiuc.edu/Research/namd/) according to best practices with no special modifications.

## Acknowledgments

The authors would like to thank Dr. Pamela Bjorkman for insightful discussions on ectodomain structures and modeling, Dr. Elisa Fadda and Dr. Mia A. Rosenfeld for their valuable discussion on Env glycosylation, and Dr. Walther Mothes for valuable discussion on Env dynamics. The authors also extend their gratitude to Dr. Abigail Dommer for assistance with membrane composition. Simulations were performed using computing resources provided by the Triton Shared Computing Cluster (TSCC) at the San Diego Supercomputer Center (SDSC). We acknowledge the use of the University of California, San Diego (UC San Diego) cryo-EM facility, which was built and equipped with funds from UC San Diego and an initial gift from the Agouron Institute. This research was supported by the National Institute of Allergy and Infectious Diseases (NIAID) of the National Institutes of Health (NIH) under Grant No. R01AI179188 and Grant No. P50-AI150464 for the Center for the Structural Biology of Cellular Host Elements in Egress, Trafficking, and Assembly of HIV (CHEETAH). M.D. is supported by a NIH PiBS T32 training grant (GM13335). S.C is a Fellow of The Jane Coffin Childs Fund for Medical Research. E.V. is an investigator of the Howard Hughes Medical Institute.

## Author Contributions

M.S. and L.C. contributed equally to this work. M.S. and L.C. prepared Env simulation models, designed and performed MD simulations, designed and performed analyses of MD simulations, and created figures and movies. M.D. and A.F. prepared the samples. M.D. acquired cryo-ET data. M.D. and S.C. analyzed cryo-ET data. E.V. oversaw cryo-ET experiments. L.C. and R.E.A. oversaw the project. M.S. and L.C. wrote the original draft of the manuscript, and all the authors contributed to reviewing and editing.

## Notes

### Competing Interest Statement

The authors have declared no competing interest.

