## Supplementary Information for "N-Glycans Modulate HIV-1 Env Conformational Plasticity"

### Table of Contents

1. **Methods**
    1.1. Computational Methods
        1.1.1. Full-length, fully glycosylated HIV-1 Env model preparation
        1.1.2. Molecular dynamics (MD) simulations
        1.1.3. Analysis of MD simulations
    1.2. Expression and cloning of enveloped virus-like particles (eVLPs)
    1.3. Cryo-electron tomography sample preparation and data collection
    1.4. Cryo-electron tomography data analysis
2. **Supplementary Figures 1 – 8**
3. **Supplementary Tables 1 – 7**
4. **Captions for Supplementary Movie 1**
5. **Supplementary References**

### 1. Methods

#### 1.1. Computational Methods

##### 1.1.1. Full-length, fully glycosylated HIV-1 Env model preparation

The human immunodeficiency virus type 1 (HIV-1) envelope glycoprotein (Env) consists of four different regions: the ectodomain, the membrane-proximal external region (MPER), transmembrane domain (TMD) and cytoplasmic tail (CT). Obtaining a structure of the full-length Env remains challenging due to the inherent structural complexity, flexibility, and extensive glycosylation. For instance, owing to its instability, the ectodomain is solved only by utilizing stabilized soluble native-like trimers known as SOSIP trimers. These trimers are stabilized by a disulfide bond between gp120 and gp41 subunits of the ectodomain, referred to as 'SOS,' and an isoleucine-to-proline point mutation, denoted as 'IP,' at residue 559[1]. SOSIP trimers are also engineered with the deletion of the furin cleavage site, a modification that prevents premature cleavage during protein expression and contributes to preserving the desired prefusion trimeric conformation, essential for structural and immunogenic studies[1]. Despite the availability of numerous high-resolution SOSIP structures, several highly flexible regions, such as loops and glycans beyond the first three monosaccharides, remain generally unresolved. Moreover, obtaining a high-resolution structure of the MPER, TMD, and CT poses considerable challenges due to their dynamic nature within the viral membrane.[2] Hitherto, only a few NMR models are available for these regions[3–5].

Here, a multi-step modeling protocol was carried out to build a full-length model of the Env as described below.

*Ectodomain modeling*

The ectodomain model (HXB2: residue 31 to 662) was built utilizing the crystal structure of BG505 SOSIP.664 trimer solved at 3.5 Å (PDB ID: 5T3Z)[6]. The SOSIP mutations and the furin cleavage site were reversed to match the gene encoding the clade A HIV-1 Env (UniProt accession number: Q2N0S6). The missing gaps of the structure were modeled as disordered loops using Modeller 10.3[7]. A total of 100 independent models of the gp120 and gp41 subunits were generated. The top models, determined based on their scoring metrics, were selected to construct multiple gp160 monomers. Specifically, the gp120-gp41 monomers were reassembled into gp160 monomers and symmetrically combined with each other to generate several models of the ectodomain trimer. These models were further visually inspected, and the model exhibiting the best structural features was selected. Particular attention was given to ensuring that no loops were entangled in knots or clashed with other parts of the structure, ensuring optimal structural integrity.

*MPER, TMD and CT modeling*

The MPER, TMD, and CT regions were modeled based on the NMR structure of the entire MPER-TMD-CT region of HIV-1 clade D (PDB ID: 7LOI)[8]. Given that a complete NMR model is not available for clade A, which is the focus of this study, we used i-TASSER to perform homology modeling using the clade D structure as a template and incorporating the clade A sequence[9,10]. Five models of confidence scores of -0.50, -1.04, -1.15, -2.07, and -4.24 were generated from i-TASSER. C-score is typically in the range of [-5,2], where a C-score of higher value signifies a model with high confidence and vice-versa. Model 1, which showed the highest confidence score, was selected. Two cysteines located in the CT of the resulting construct (C764 and C837) were palmitoylated using the lipid-tail functionalization widget available within *Glycan Reader* in CHARMM-GUI[11].

*Full-length model assembly*

Since the ectodomain model and the MPER-TMD-CT model were constructed from two independently resolved structures, their direct connection was hindered by misalignment and structural distortions, necessitating additional modeling to achieve an accurate and seamless integration. To address this, we first re-modeled the ectodomain's HR2 helix up to residue 668 (namely residues 637–668, which encompass part of the MPER's C-terminal helix, 660–668) using AlphaFold2[12] and removed the unfolded N-terminal residues (663–668) from the MPER-TMD-CT model. Then, via the re-modeled HR2 helix, the ectodomain was seamlessly connected to the MPER-TMD-CT model, ensuring proper alignment and structural integrity. The connection points were refined through local energy minimization with the autoIMD plugin in VMD.[13] Superimposition of the MPER region in our model with the corresponding MPER region resolved in the crystal structure (PDB ID: 4G6F[14]) revealed remarkable structural similarity, underscoring the high quality of our model (see Fig. S7).

*Glycosylation*

The HIV-1 Env glycoprotein harbors 23 N-glycan (N-X-S/T) sites per monomer, which have been shown to be heterogeneously populated across various glycoanalytic studies[15–17]. Therefore, for the scope of this study, a heterogenic (i.e., not uniform across monomers) site-specific glycoprofile has been derived based on the available glycoanalytic data[15–17]. A detailed, per-chain description of the site-specific glycoprofile for the fully glycosylated model of the HIV-1 Env glycoprotein simulated in this work is provided in Tables S1–S3. Glycans were modeled using the *Glycan Reader & Modeler* tool[18] available in CHARMM-GUI[11]. We note that the oligosaccharides (GlcNAc-/GlcNAc2-/ManGlcNAc2-) originally solved in the cryo-EM structures have generally been preserved and used as a scaffold wherever feasible for constructing complete glycans. However, moderate adjustments to glycan dihedrals and/or asparagine side chains have been carried out at densely glycosylated sites to eliminate steric clashes and accommodate all the glycans.

*Membrane building*

The lipid composition of the membrane patch was selected based on the experimental studies of the HIV-1 plasma membrane compositions[19–22]. The outer and inner leaflet abundances were derived from previous molecular dynamics (MD) simulations and lipidomic data of mammalian cell plasma membranes[23] (Table S4). An asymmetric lipid membrane patch (200 Å × 200 Å) was constructed using CHARMM-GUI's input generator[11,24]. The ratio of short-chain/saturated to long-chain/unsaturated phosphatidylcholine (PC) lipids was set to 40:60[20]. These fractions were adjusted to compensate for the simplification of the membrane, which artificially increased the lipid count of the inner leaflet due to bilayer asymmetry.

*System preparation*

After functionalization (i.e., glycosylation and palmitoylation) of the HIV-1 Env glycoprotein, protonation states were assigned using PROPKA3[25] at pH 7.4. The generated models were parameterized using *psfgen* and the CHARMM36m all-atom additive force fields for proteins, lipids, and glycans[26–28]. The systems were solvated with TIP3P[29] explicit water model in a box, ensuring a minimum distance of 24 Å between the protein and the box edges. $Na^+$ and $Cl^-$ ions were added at a concentration of 150 mM were added to neutralize the charge of the system. Solvation and charge neutralization were performed using the *solvate* and *autoionize* plugins within VMD[13].

#### 1.1.2. Molecular dynamics (MD) simulations

All-atom MD simulations were performed on the Triton Shared Computing Cluster (TSCC) at the San Diego Supercomputer Center (SDCS) using NAMD3[30]. A multi-step approach involving repeated minimization cycles, NVT melting (for the membrane), and NPT equilibration was applied to relax the systems. Initially, the systems were subjected to an initial 10,000 energy minimization steps using the conjugate gradient energy method during which the protein, glycans, lipid heads (P atom for DPPC, POPC, POPE, PSM and POPS, and O3 atom for CHL), solvent, and ions were kept fixed. Next, to allow equilibration of the lipid tails (i.e., melting), the temperature was gradually increased from 10 K to 310 K over 0.5 ns (NVT ensemble) using a time step of 1 fs. This was followed by a further 20,000 steps of energy minimization, during which positional harmonic restraints with a force constant of 1.0 (kcal/mol)/$Å^2$ were applied to the protein and glycan atoms. A 0.5 ns NPT (isothermal-isobaric) equilibration was then conducted at 2 fs/step using an anisotropic pressure coupling scheme (*useFlexibleCell yes, useConstantArea no*) with the harmonic restraints on protein and glycan retained. Next, all restraints were released, and another 0.5 ns NPT (isothermal-isobaric) equilibration was performed. Lastly, the pressure coupling was switched to a semi-isotropic scheme (*useFlexibleCell yes, useConstantArea yes*), and another 20 ns of NPT equilibration was conducted. Prior to production MD simulations, all the systems were meticulously inspected to ensure no structural artifacts were present. Five replicates of NPT production MD simulations with semi-isotropic pressure coupling were performed for 1.05 μs. It is important to note that all replicates were conducted independently, from their respective minimization and continuing through to the production run. All simulations were performed with an integration time step of 2 fs, utilizing the SHAKE[31] algorithm to constrain covalent bonds involving all hydrogen atoms. Periodic boundary conditions were employed during the simulations, with the particle-mesh Ewald[32] method used for long-range electrostatics calculations. A maximum grid spacing of 2 Å was applied. Non-bonded interactions, including van der Waals forces and short-range electrostatics, were handled using a cutoff distance of 12 Å. Control of the temperature and pressure during the simulations was achieved with a Langevin thermostat[33] (310 K) and a Nosé-Hoover Langevin barostat[34,35] (1.01325 bar) as implemented in NAMD. A summary of the simulations performed in this work is provided in Table S5.

#### 1.1.3. Analysis of MD simulations

*Tilting angle calculation*

The tilting angle of the Env protein was calculated using an in-house Tcl script executed within VMD[13]. For this calculation, two vectors lying in the same plane, perpendicular to the membrane, were defined. Specifically, a vector representing the center of mass (COM) of the protein core, specifically HR1C residues 572–597, and a vector representing the principal axis normal to the membrane were computed at each simulation frame. The tilting angle was determined by measuring the angle between these two vectors using dot products and trigonometric calculations. Each simulation frame was analyzed to determine the Env's tilting angle at that frame. According to the value of the Env's tilting angle, each frame was then assigned to the ensemble of conformations accounting for the range of values that included that angle (e.g., either 0–10°, 10–20°, 20–30°, or 30°+). At the end of this process, the four ensembles representing the four tilting angle ranges were generated and used for subsequent comprehensive collective analysis.

*Glycans-membrane contacts*

Glycan-membrane interactions were analyzed using the *measure contacts* command in VMD,[13] supplemented by custom made Tcl scripts. Contacts were defined using a 4.5 Å cutoff distance between glycan and membrane heavy atoms on a residue basis. For each glycan, the total number of frames where contacts occurred was determined, and the contact frequency was calculated as a percentage of frames in which interactions were observed relative to the total simulation frames. For the N88 and N611 glycans, the time evolution of their contacts with the membrane was analyzed, along with an assessment of the cooperativity of their interactions. This analysis evaluated the simultaneous interactions of multiple glycans, either originating from the same site (e.g., two N88 or two N611) or from different sites (e.g., one N88 and one N611).

*Conservation analysis of Env*

To analyze the evolutionary conservation of gp120 and gp41, we used the Evolutionary Conservation Profiles of Proteins server (Consurf)[36–39]. Homologues were retrieved from the UniProt Reference Clusters database at 90% sequence identity (UNIREF90) using the HMMER algorithm (HMMER) with an E-value cutoff of 0.0001 and a single iteration. The homologous sequences were refined by clustering at 95% sequence identity using the Cluster Database at High Identity with Tolerance algorithm (CD-HIT), retaining the highest-scoring sequence when overlaps exceeded 10%. Homologues with less than 60% coverage of the query sequence were excluded, and up to 300 sequences were sampled for further analysis. The multiple sequence alignments were generated using the Multiple Alignment using Fast Fourier Transform method (MAFFT). A phylogenetic tree was constructed using the Neighbor Joining method (NJ) with maximum likelihood distances. Conservation scores were calculated using a Bayesian approach with the most appropriate amino acid substitution model determined by best fit.

*MPER and TMD hinge angles* calculation

The MPER and TMD hinge angles were calculated using a custom Tcl script executed within VMD[13]. For the MPER hinge angle, three points lying in the same plane were defined: the alpha carbon (Cα) of residue 624, the COM of residues 635–639's Cα, and the Cα of residue 651. Similarly, for the TMD hinge angle, three points were defined: the Cα of residue 630, the Cα of residue 651, and the Cα of residue 664. The angles were computed for each frame of the MD trajectories by measuring the angle formed by the three defined points, with trigonometric calculations based on the dot product of vectors used to derive the values.

*TMD inclination angles calculation*

To calculate the TMD inclination angle was calculated using a custom Tcl script executed within VMD[13]. For this angle, we defined two vectors lying in the same plane, perpendicular to the membrane. The first vector was aligned along a segment of the TMD defined by the Cα atoms of residues 684–696. The second vector was defined as parallel to the membrane plane. The inclination angle was determined for each frame of the MD trajectories by measuring the angle between these two vectors, using trigonometric calculations based on the dot product.

*Accessible surface area (ASA)*

ASA was calculated using the *measure sasa* command in VMD,[13] which utilizes the Shrake and Rupley algorithm,[40] in combination custom Tcl scripts. The ASA was computed using probe radii ranging from 1.4 Å to 8 Å. The glycan-shielded area was determined by subtracting the ASA of the *protein-with-glycans* from the ASA of the *protein-without-glycans*. Similarly, the membrane-occluded area was calculated by subtracting the ASA of the *protein-with-glycans-and-membrane* from the ASA of the *protein-with-glycans*. ASA values were computed at 1-nanosecond intervals throughout the simulations, The values were averaged across all the respective replicas and standard deviation was computed.

*Epitope-specific accessible surface area (ASA)*

The ASA of key antigenic regions of the Env trimer including the (MPER) (residues 628–651) the gp41-gp120 interface (residues 4–19, 46–56, 59–64, 207–209, 503–511, 580–584, 586–593, 596–606), the CD4 binding site (CD4bs) (residues 333–341, 397–400, 423–430, 435–443), the V1V2 loop (residues 94–168), the V3 loop (residues 266–299), the fusion peptide (FP) (residues 480–494), and the silent face epitope (residues 303–306, 413–416, 260–263, 222, 183–184, 27–29) was computed,. We note that the residue numbering provided here corresponds to our Env model, which may differ from the standard HXB2 reference sequence. For clarity and accurate mapping, refer to Figure S1 for alignment of our model with the HXB2 sequence, allowing easy conversion between the two numbering schemes. Additionally, the supplementary PDB/DCD files can be used for direct visualization of these epitopes. ASA values were computed at 1-nanosecond intervals across all simulations following the same methodology described in the previous section. The analysis was performed using a 7.2 Å probe radius**,** approximating the size of the complementarity-determining region (CDR) of an antibody's variable fragment (Fv). The values were averaged across all the respective replicas and standard deviation was computed.

*MPER peptide accessible surface area (ASA)*

To evaluate the accessibility of the MPER peptide (residue 671–683), the trajectory frames were clustered into four distinct ensembles based on the degree of Env tilting observed (i.e., 0–10°, 10–20°, 20–30°, and 30°+). These ensembles were then used to calculate the ASA as a function of the Env's tilting angle, providing insights into how tilting affects the epitope accessibility. For every frame within each ensemble, ASA values were separately computed for each chain using a 7.2 Å probe radius**,** approximating the size of the complementarity-determining region (CDR) of an antibody's variable fragment (Fv). The highest ASA values were then extracted to identify the most accessible chain. ASA values were averaged across each ensemble. The glycan-shielded and membrane-occluded areas were calculated for each tilt angle range as previously described. To assess the uncertainty of these means, bootstrap resampling with 2000 iterations was conducted, providing an estimation of the standard error of the mean.

### 1.2. Expression and cloning of enveloped virus-like particles (eVLP)

*EVLP cloning*

The full-length envelope glycoprotein sequence for BG505 was obtained from UniProt (ABA61515.1) and reverse transcribed using GenScript's codon optimization tool and optimized for human expression. GenScript cloned this fragment into the pcDNA3.1(+) backbone and upon receipt, this plasmid was expanded by transformation in Stbl3 competent cells. To clone in the signal peptide of gp160, the second BstEII restriction site was mutated, and the peptide sequence was added by Gibson assembly. To improve Env expression in the eVLPs, an endocytosis prevention motif (EPM) and an ESCRT and ALIX-binding region (EABR) sequence were added to the c-terminal side of the Env sequence. This technology is described in Hoffman et al., 2023.

*eVLP expression and purification*

293FT cells were grown to 80-90% confluency and fresh media was added just prior to transfection of the eVLP plasmid using FuGENE transfection reagent. The transfected cells were incubated at 37°C and 5% $CO_2$ for 72 hours. After 72 hours, the supernatant was collected and centrifuged at 400 × g for 10 minutes. The supernatant was filtered through a 0.45 µm syringe to remove excess cell debris that remained after centrifugation. Supernatant was transferred to a 100 kDa spin column and centrifuged up to 4000 × g to reduce the total volume of the sample to less than 5 mL. The concentrated sample was layered on top of a 2 mL 20% sucrose cushion in an ultracentrifuge tube. The sample was spun at 135,000 × g for 2 hours at 4°C. The supernatant was carefully removed and the pellet re-suspended in 1xPBS. The re-suspended eVLPs were incubated in the ultracentrifuge tube overnight at 4°C. The following morning, the eVLPs were carefully mixed and pipetted into a 100k MWCO spin column. To remove excess sucrose contamination, the sample was rinsed 3 times with 1xPBS by spinning at 10,000 × g at 4°C in the spin column. After the final rinse, the column was spun to reduce the final volume of the sample to 20-50 µL. This purification process was originally performed in T75 flasks but was ultimately scaled up to 15 mm dishes to increase throughput.

### 1.3. Cryo-electron tomography sample preparation and data collection

*Cryo-EM grid preparation of eVLPs*

Before sample deposition, gold QUANTIFOIL R 2/1 cryo-EM grids were plasma-cleaned for 60 seconds using a Pelco plasma cleaner equipped with air at 25W power. 4 µL of purified eVLPs was applied to the grids and back-blotted using the Leica EM GP2 Automatic Plunge Freezer and then plunge frozen into liquid ethane or a 50/50 ethane propane mix. The blot time varied from 4 – 10 seconds, depending on the viscosity of the eVLP prep. Multiple blot times would be used for each eVLP sample to determine the best conditions for that sample preparation.

*Cryo-ET data collection*

For all datasets collected, a Titan Krios-3 TEM (Thermo Fisher) operating at 300 kV with K3 direct electron detector (Gatan) at a nominal magnification of 64,000x (pixel size 1.341 Å, counting mode) was used. A 70µm objective aperture was inserted. A Gatan zero-loss imaging filter with a 15 eV-wide slit was used throughout the data collection. The defocus range was -3 to -5 µm. paceTOMO in serialEM[41] was used for collecting tilt series automatically. Targets were selected manually to maximize the number of eVLPs that could collected within each tilt series. For each tilt series, a dose-symmetric scheme was applied with a tilt range of -54° to +54° with an increment of 3°, resulting in 37 tilt images being collected with 8 frames in each tilt. The total dose was 140 e-/Å$^2$ and the dose rate was set at 15.9 e$^-$/Å$^2$/s. The protocol describing detailed steps in data collection is available online [42].

#### 1.4. Cryo-electron tomography data analysis

*Tilt series alignment and tomogram reconstruction*

Linux Warp or Windows Warp[43] was used for the pre-processing of tomograms before subtomogram analysis. Specifically, motion correction, CTF estimation and gain correction were performed in Warp and then all tilt series were aligned with either areTomo, or eTomo in IMOD[44,45] v4.11.25 using patch tracking. The Residual error mean (in nm) calculated from eTomo was used to assess the alignment quality for different tilt series. Tomograms with <2 nm Residual error mean are eligible for future analysis. Next, the alignment data was fed back to Warp for tomogram reconstruction using weighted back-projection at 10 Å/pxl. Tomogram visualization and Env particle picking were performed on deconvoluted tomograms generated from Warp with default settings.

*Env particle picking and initial orientation assignment in Dynamo*

IMOD was used to visualize all reconstructed tomograms and manually pick Env subtomograms from VLPs. In total, 705 tomograms were inspected and 4,743 particles were picked. Picking for each VLP was conducted by first placing a point in the center of the VLP using the XYZ view in IMOD as a guide to determine the center in 3-dimesional space, placing a second point on the membrane of the VLP, and then finally points were added for each Env that could be visually identified. The coordinates of selected Envs from each VLP were analyzed in Dynamo in matlab[46–49] v11509. VLPs are modeled as spheres with manually defined center and radius in Dynamo. The initial orientation of each Env subtomogram are assigned perpendicular to the surface of the VLP model it belongs to. The azimuth angle was randomized to minimize the effect of the missing wedge. The coordinates were then used for further subtomogram analysis in Relion3[50].

*Subtomogram analysis and three-dimensional model reconstruction*

For the 3D reconstruction of Env, a total of 4,743 subtomograms were combined and provided to Relion3 following particle extraction with Warp. These included WT subtomograms as well as N611A and N88A mutant forms, which were originally generated for future functional studies but were also used here to supplement the dataset and increase particle numbers. The mutant datasets were not analyzed independently and were used solely to improve alignments. No mutant-specific analyses are presented in this study.

The subtomograms were binned to 10 Å/pxl at this stage. Subtomograms were refined in Relion3 with C1 symmetry first to align the membrane, then followed by a 6-class 3D classification with ang- and psi-angle limit to local ranges. To avoid over-fitting, the regularization parameter T was set to 1 instead of default 4 and the resolution E-step was set to 25Å to disable alignment of high-frequency signals. Good class-averages were chosen for further C1 alignment. The final map reported in this study only contained WT particles and had a resolution of 33Å as determined by the gold-standard Fourier shell correlation (FSC) using the 0.143 criterion.

UCSF ChimeraX[51] v1.9 was used for all the volume segmentation, figure and movie generation, and automatic rigid-body docking processes.

*Geometric analysis of Env tilt angle relative to VLP surface*

After obtaining the Env reconstruction, subtomograms included in the final map were used to calculate their orientation relative to VLP. Envs were assessed for tilting on a per-particle basis. The angle between the Env orientation vector (determined through subtomogram averaging) and a vector perpendicular to the membrane at each respective Env position was determined as the Env tilting angle. To calculate the vector perpendicular to the membrane, focused angular refinement on the membrane region was performed, resulting in a disordered Env ectodomain and a well-defined membrane density. Tilt angle distributions were plotted as a histogram, with 0° indicating no tilt and 90° indicating an orientation parallel to the VLP membrane. A total of seven outlier points were removed using the ROUT method with $Q = 1\%$. The outliers all had values >90°, indicating that they were likely misaligned particles.

### 2. Supplementary Figures

```
HXB2                                     MRVKEKYQHLWRWGWRWGTMLLGMLMICSATEKLWVTVYYGVPVWKEATTTLFCASDAKA      60
Full_legnth_env_based_on_uniprot_Q2N0S6  ------------------------------AENLWVTVYYGVPVWKDAETTLFCASDAKA      30
                                                                       :*:*************:* ***********

HXB2                                     YDTEVHNVWATHACVPTDPNPQEVVLVNVTENFNMWKNDMVEQMHEDIISLWDQSLKPCV      120
Full_legnth_env_based_on_uniprot_Q2N0S6  YETEKHNVWATHACVPTDPNPQEIHLENVTEEFNMWKNNMVEQMHTDIISLWDQSLKPCV      90
                                         *:** ******************: * ****:******:****** **************

HXB2                                     KLTPLCVSLKCTDLKNDTNTNSSSGRMIMEKGEIKNCSFNISTSIRGKVQKEYAFFYKLD      180
Full_legnth_env_based_on_uniprot_Q2N0S6  KLTPLCVTLQCTNVTNNITDD--------MRGELKNCSFNMTTELRDKKQKVYSLFYRLD      142
                                         *******:*:**::.*: . :         :**:******::*.:*.* ** *::**:**

HXB2                                     IIPIDN--------DTTSYKLTSCNTSVITQACPKVSFEPIPIHYCAPAGFAILKCNNKT      232
Full_legnth_env_based_on_uniprot_Q2N0S6  VVQINENQGNRSNNSNKEYRLINCNTSAITQACPKVSFEPIPIHYCAPAGFAILKCKDKK      202
                                         :: *::        ....*:* .****.****************************::*.

HXB2                                     FNGTGPCTNVSTVQCTHGIRPVVSTQLLLNGSLAEEEVVIRSVNFTDNAKTIIVQLNTSV      292
Full_legnth_env_based_on_uniprot_Q2N0S6  FNGTGPCPSVSTVQCTHGIKPVVSTQLLLNGSLAEEEVMIRSENITNNAKNILVQFNTPV      262
                                         ******* .**********:******************:*** *:*:***.*:**:** *

HXB2                                     EINCTRPNNNTRKRIRIQRGPGRAFVTIG-KIGNMRQAHCNISRAKWNNTLKQIASKLRE      351
Full_legnth_env_based_on_uniprot_Q2N0S6  QINCTRPNNNTRKSIRI--GPGQAFYATGDIIGDIRQAHCTVSKATWNETLGKVVKQLRK      320
                                         :************ ***  ***:** : *  **::*****.:*:*.**:** ::..:**:

HXB2                                     QFGNNKTIIFKQSSGGDPEIVTHSFNCGGEFFYCNSTQLFNSTWFNSTWSTEGSNNTEGS      411
Full_legnth_env_based_on_uniprot_Q2N0S6  HFGNNTIIRFANSSGGDLEVTTHSFNCGGEFFYCNTSGLFNSTWISNT-SVQGSNSTGSN      379
                                         :****. * * :***** *:.**************:: ******:..* *.:***.* ..

HXB2                                     DTITLPCRIKQIINMWQKVGKAMYAPPISGQIRCSSNITGLLLTRDGGNSNNESEIFRPG      471
Full_legnth_env_based_on_uniprot_Q2N0S6  DSITLPCRIKQIINMWQRIGQAMYAPPIQGVIRCVSNITGLILTRDGGSTNSTTETFRPG      439
                                         *:***************::*:*******.* *** ******:******.:*. :* ****

HXB2                                     GGDMRDNWRSELYKYKVVKIEPLGVAPTKAKRRVVQREKRAVGIGALFLGFLGAAGSTMG      531
Full_legnth_env_based_on_uniprot_Q2N0S6  GGDMRDNWRSELYKYKVVKIEPLGVAPTRAKRRVVGREKRAVGIGAVFLGFLGAAGSTMG      499
                                         ****************************:****** **********:*************

HXB2                                     AASMTLTVQARQLLSGIVQQQNNLLRAIEAQQHLLQLTVWGIKQLQARILAVERYLKDQQ      591
Full_legnth_env_based_on_uniprot_Q2N0S6  AASMTLTVQARNLLSGIVQQQSNLLRAIEAQQHLLKLTVWGIKQLQARVLAVERYLRDQQ      559
                                         ***********:*********.*************:************:*******:***

HXB2                                     LLGIWGCSGKLICTTAVPWNASWSNKSLEQIWNHTTWMEWDREINNYTSLIHSLIEESQN      651
Full_legnth_env_based_on_uniprot_Q2N0S6  LLGIWGCSGKLICTTNVPWNSSWSNRNLSEIWDNMTWLQWDKEISNYTQIIYGLLEESQN      619
                                         *************** ****:****:.*.:**:: **::**:**.***.:*:.*:*****

HXB2                                     QQEKNEQELLELDKWASLWNWFNITNWLWYIKLFIMIVGGLVGLRIVFAVLSIVNRVRQG      711
Full_legnth_env_based_on_uniprot_Q2N0S6  QQEKNEQDLLALDKWASLWNWFDISNWLWYIKIFIMIVGGLIGLRIVFAVLSVIHRVRQG      679
                                         *******:** ***********:*:*******:********:**********:::*****

HXB2                                     YSPLSFQTHLPTPRGPDRPEGIEEEGGERDRDRSIRLVNGSLALIWDDLRSLCLFSYHRL      771
Full_legnth_env_based_on_uniprot_Q2N0S6  YSPLSFQTHTPNPRGLDRPERIEEEDGEQDRGRSTRLVSGFLALAWDDLRSLCLFCYHRL      739
                                         ********* *.*** **** ****.**:**.** ***.* *** **********.****

HXB2                                     RDLLLIVTRIVELLGR-------RGWEALKYWWNLLQYWSQELKNSAVSLLNATAIAVAE      824
Full_legnth_env_based_on_uniprot_Q2N0S6  RDFILIAARIVELLGHSSLKGLRLGWEGLKYLWNLLAYWGRELKISAINLFDTIAIAVAG      799
                                         **::**.:*******:        ***.*** **** **.:*** **:.*::: *****

HXB2                                     GTDRVIEVVQGACRAIRHIPRRIRQGLERILL   856
Full_legnth_env_based_on_uniprot_Q2N0S6  WTDRVIEIGQRLCRAFLHIPRRIRQGFERALL   831
                                          ******: *  ***: *********:** **
```

**Supplementary Figure 1.** Sequence alignment of the full-length Env model used in this study with the reference HXB2 Env sequence.

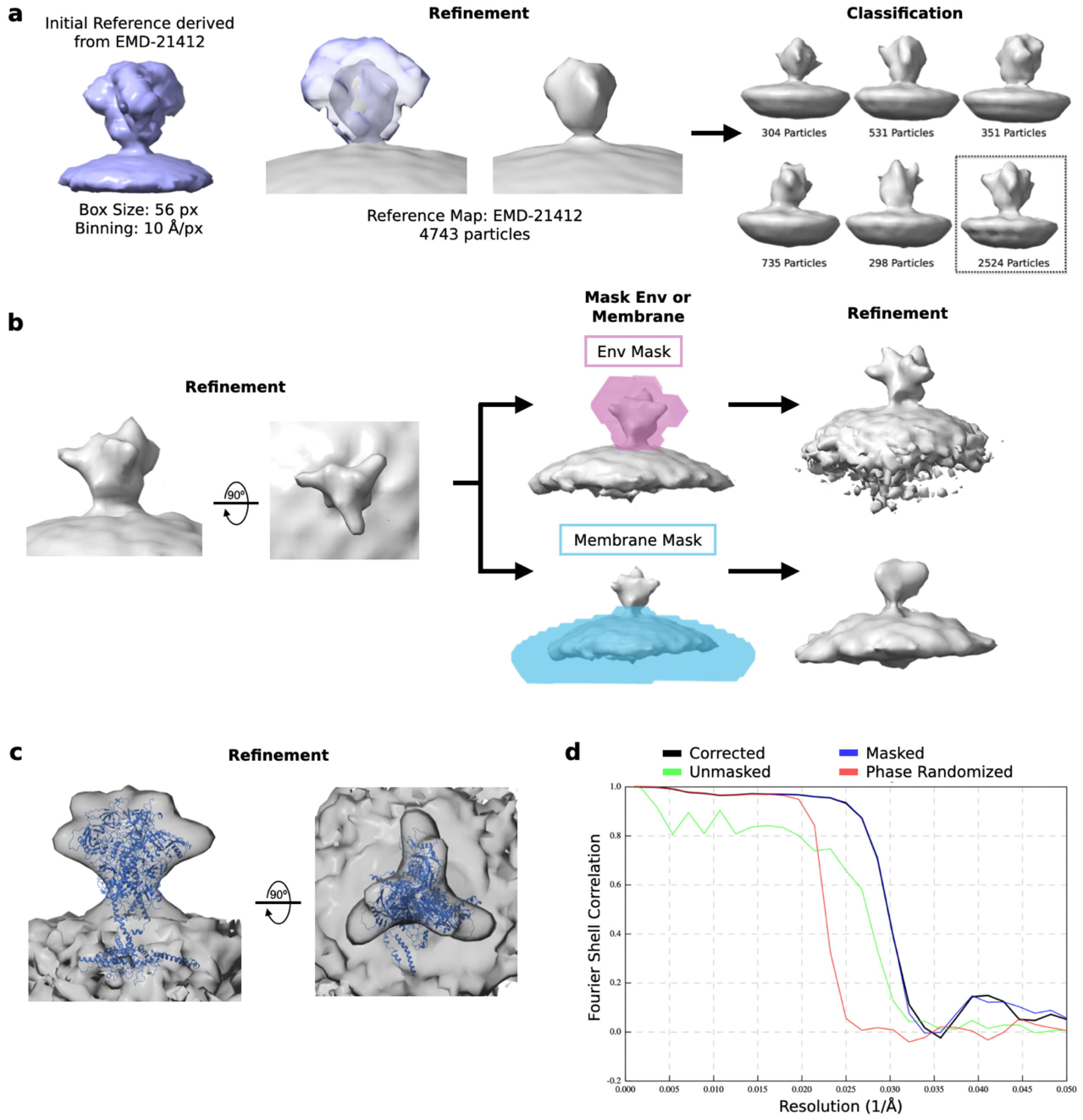


**Supplementary Figure 2.** (**a**) The initial reference used for refinement in Relion3 was derived from EMD-21412 to roughly align the Env particles. After initial refinement, the particles were classified into six classes to remove junk particles and the most reasonable class was selected for further refinement, as indicated by the box. (**b**) The selected class was mask refined with an Env mask and a membrane mask to generate two separate refinements with an optimally aligned Env or membrane that is used for downstream angle determination. (**c**) Final reconstruction of WT Env with the model from this paper fitted into the density. (**d**) Gold-standard Fourier shell correlation (FSC) curve using the 0.143 criterion. Estimated resolution of 33 Å.

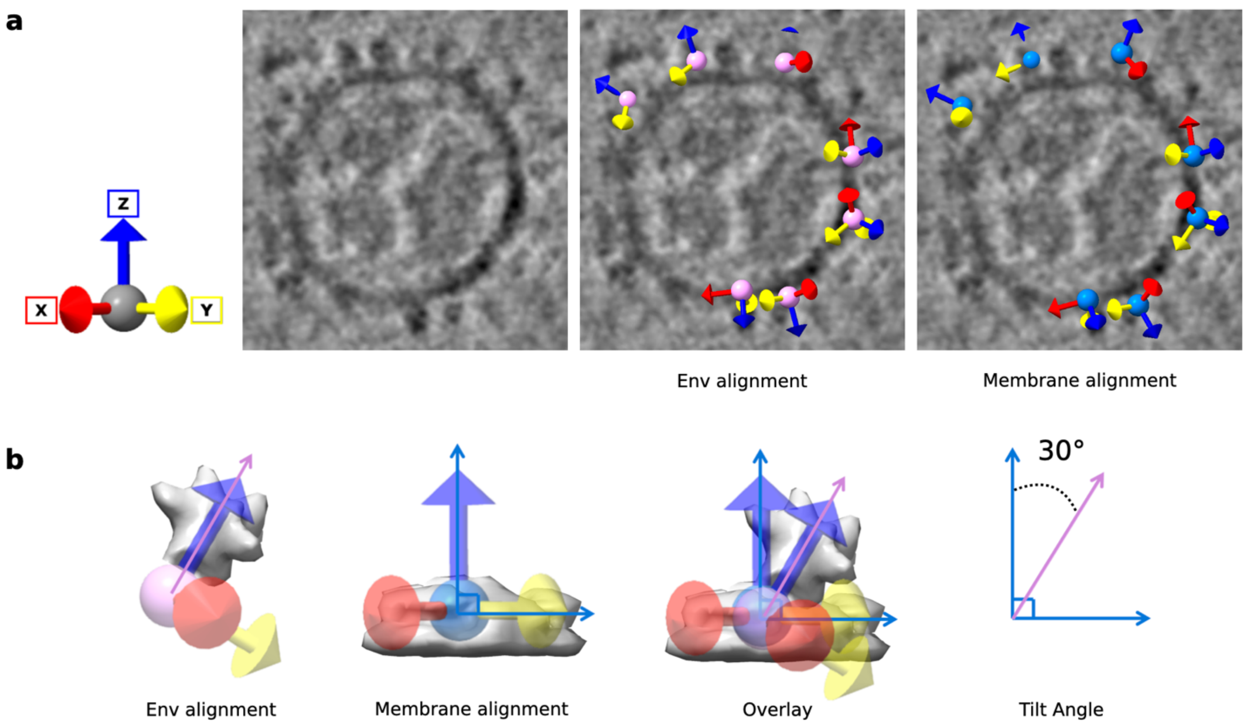


**Supplementary Figure 3.** (**a**) Particle orientation is depicted with the blue arrow indicating the z-axis, red arrow indicating the x-axis, and the yellow arrow indicating the y-axis. Left: Example tomogram slice shown. Middle: Env-masked and aligned particles, indicated by the pink sphere, overlayed on tomogram slice. Right: Membrane-masked and aligned particles, indicated by the blue sphere, overlayed on the same tomogram slice. (**b**) Simplified schematic of the angle determination for Env tilting relative to the viral membrane based on the alignment of the masked particles.

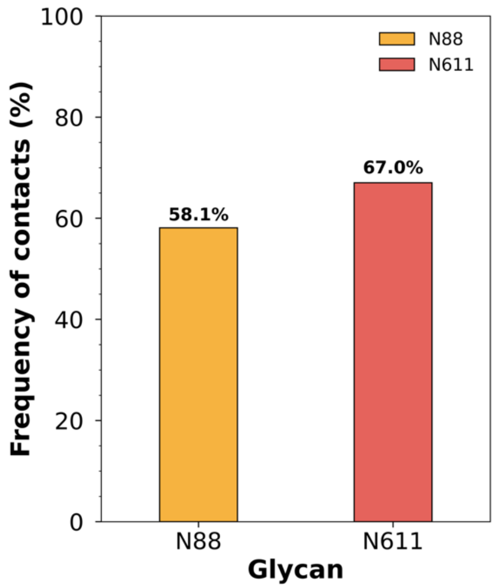


**Supplementary Figure 4.** Bar plot showing the frequency of contacts between N-glycans N88 (yellow) and N611 (red) with the lipid membrane across all simulations and protomers, expressed as % frames with at least one contact from either is observed.

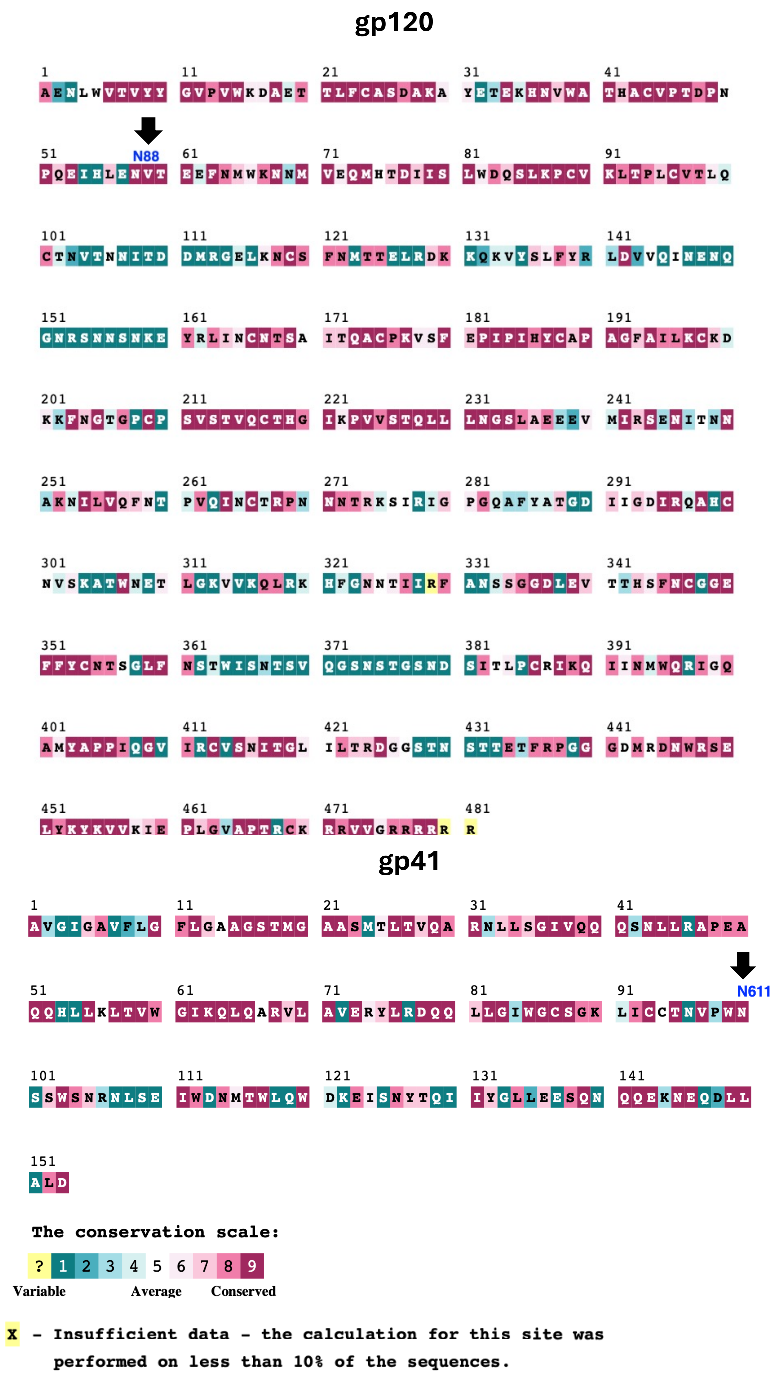

**Supplementary Figure 5.** gp120 and gp41 sequences highlighting conserved and variable residues across HIV-1 strains.

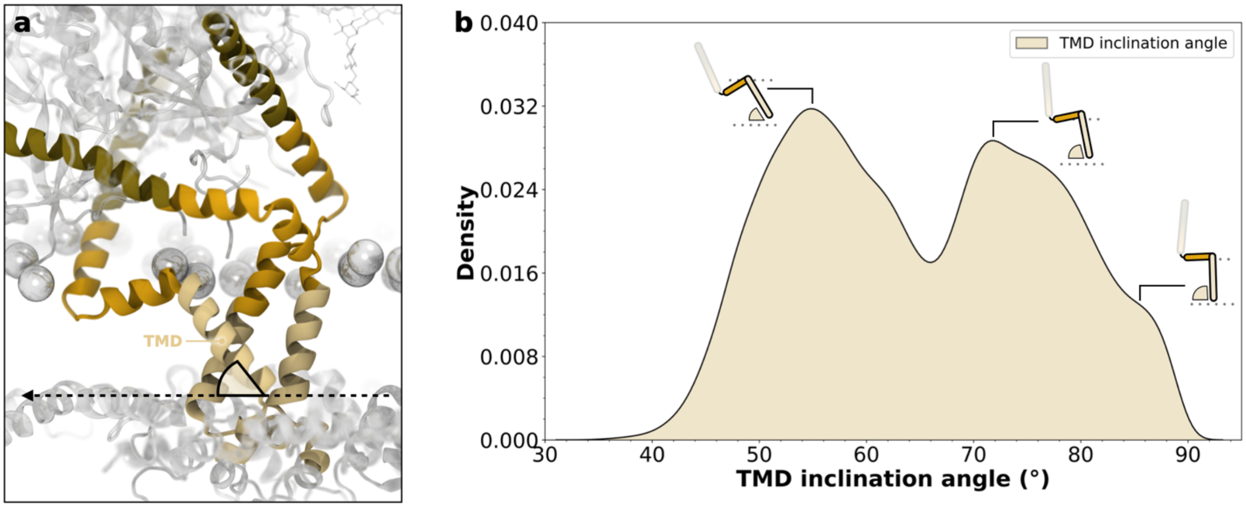


**Supplementary Figure 6.** (**a**) Structural depiction of the Env trimer's transmembrane domain (TMD) showing the inclination angle relative to the membrane plane. The TMD helices are shown in yellow. The dashed black line indicates the membrane plane, and the angle measurement is illustrated with a black triangle.
(**b**) Kernel density estimate of the TMD inclination angle distribution.

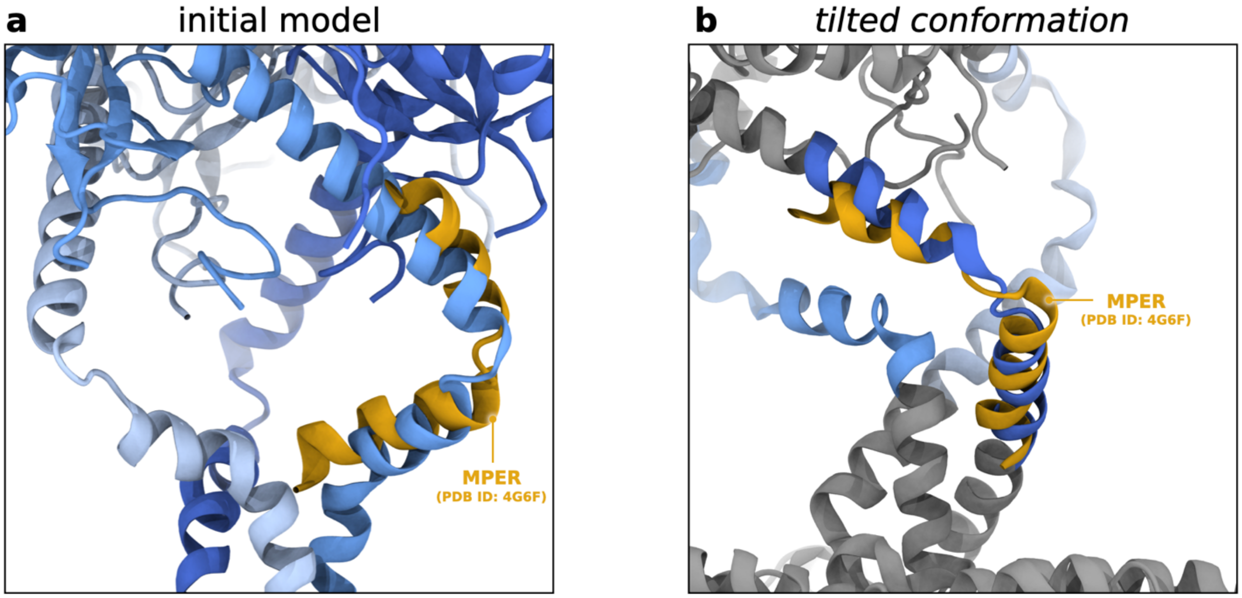


**Supplementary Figure 7.** Alignment of the MPER region from the crystal structure of 10E8 Fab in complex with the HIV-1 Env's MPER (yellow cartoons, PDB ID: 4G6F[14]) onto the HIV-1 Env's MPER as modeled in this study (blue cartoons). (**a**) Alignment onto the initial model. (**b**) Alignment onto a tilted conformation, resulting in a most exposed conformation of the MPER.

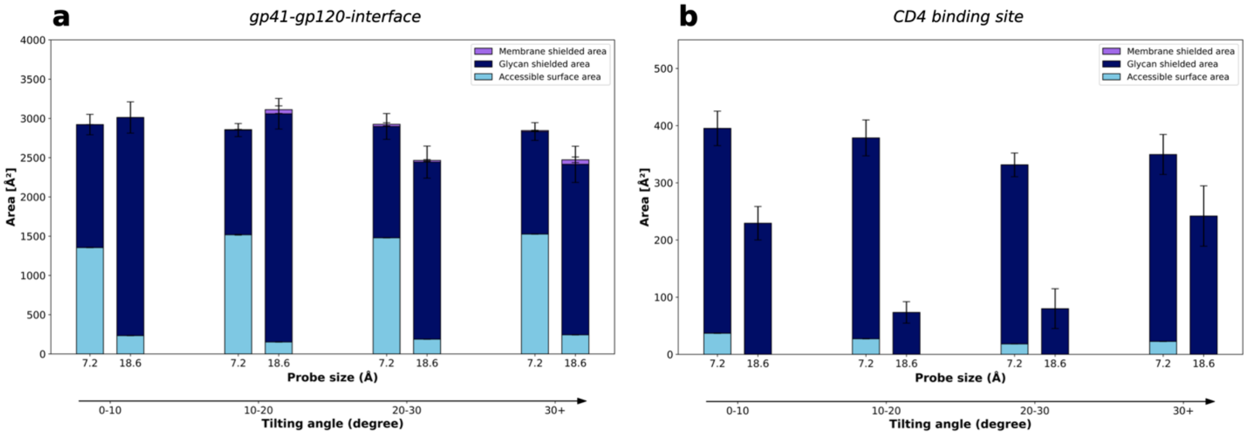


**Supplementary Figure 8.** Quantification of the membrane-shielded area (purple), glycan-shielded area (dark blue), and accessible surface area (light blue) for the gp41-gp120 interface (**a**) and the CD4 binding site (**b**) across tilting ensembles (0–10°, 10–20°, 20–30°, and 30+°) for probe sizes of 7.2 Å and 18.6 Å. Error bars represent variability across simulation replicates.

### 3. Supplementary Tables

**Supplementary Table 1.** Glycan compositions for chain A. Glycosylation sites are reported using HXB2 notation.

CHAIN A

| # | SITE | TYPE | STRUCTURE | SEQUENCE |
|---|---|---|---|---|
| G1 | N88 | FA2 | 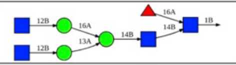 | bDGlcNAc(1→2)aDMan(1→6)[bDGlcNAc(1→2)aDMan(1→3)]bDMan(1→4)bDGlcNAc(1→4)<br>[aLFuc(1→6)]bDGlcNAc(1→)PROA-88 |
| G2 | N133 | FA2 | 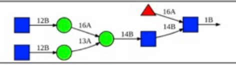 | bDGlcNAc(1→2)aDMan(1→6)[bDGlcNAc(1→2)aDMan(1→3)]bDMan(1→4)bDGlcNAc(1→4)<br>[aLFuc(1→6)]bDGlcNAc(1→)PROA-133 |
| G3 | N137 | FA2G2S2 | 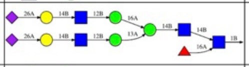 | aDNeu5Ac(2→6)bDGal(1→4)bDGlcNAc(1→2)aDMan(1→6)<br>[aDNeu5Ac(2→6)bDGal(1→4)bDGlcNAc(1→2)aDMan(1→3)]bDMan(1→4)bDGlcNAc(1→4)<br>[aLFuc(1→6)]bDGlcNAc(1→)PROA-137 |
| G4 | N156 | M9 | 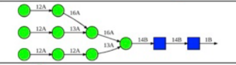 | aDMan(1→2)aDMan(1→6)[aDMan(1→2)aDMan(1→3)]aDMan(1→6)<br>[aDMan(1→2)aDMan(1→2)aDMan(1→3)]bDMan(1→4)bDGlcNAc(1→4)bDGlcNAc(1→)PROA-156 |
| G5 | N160 | M6 | 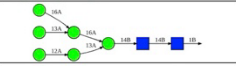 | bDGal(1→4)bDGlcNAc(1→2)aDMan(1→3)[aDMan(1→6)<br>[aDMan(1→3)]aDMan(1→6)]bDMan(1→4)bDGlcNAc(1→4)bDGlcNAc(1→)PROA-160 |
| G6 | N197 | FA2G2S1 | 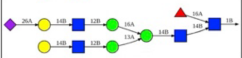 | aDNeu5Ac(2→6)bDGal(1→4)bDGlcNAc(1→2)aDMan(1→6)<br>[bDGal(1→4)bDGlcNAc(1→2)aDMan(1→3)]bDMan(1→4)bDGlcNAc(1→4)[aLFuc(1→6)]bDGlcNAc(1→)PROA-197 |
| G7 | N234 | M9 | 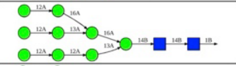 | aDMan(1→2)aDMan(1→6)[aDMan(1→2)aDMan(1→3)]aDMan(1→6)<br>[aDMan(1→2)aDMan(1→2)aDMan(1→3)]bDMan(1→4)bDGlcNAc(1→4)bDGlcNAc(1→)PROA-234 |
| G8 | N262 | M8 | 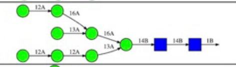 | aDMan(1→2)aDMan(1→6)[aDMan(1→3)]aDMan(1→6)<br>[aDMan(1→2)aDMan(1→2)aDMan(1→3)]bDMan(1→4)bDGlcNAc(1→4)bDGlcNAc(1→)PROA-262 |
| G9 | N276 | Hybrid G1 | 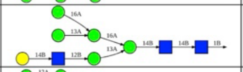 | bDGal(1→4)bDGlcNAc(1→2)aDMan(1→3)[aDMan(1→6)<br>[aDMan(1→3)]aDMan(1→6)]bDMan(1→4)bDGlcNAc(1→4)bDGlcNAc(1→)PROA-276 |
| G10 | N295 | M8 | 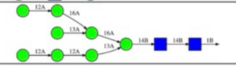 | aDMan(1→2)aDMan(1→6)[aDMan(1→3)]aDMan(1→6)<br>[aDMan(1→2)aDMan(1→2)aDMan(1→3)]bDMan(1→4)bDGlcNAc(1→4)bDGlcNAc(1→)PROA-295 |
| G11 | N301 | M9 | 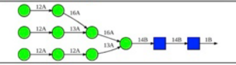 | aDMan(1→2)aDMan(1→6)[aDMan(1→2)aDMan(1→3)]aDMan(1→6)<br>[aDMan(1→2)aDMan(1→2)aDMan(1→3)]bDMan(1→4)bDGlcNAc(1→4)bDGlcNAc(1→)PROA-301 |
| G12 | N339 | M9 | 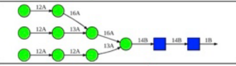 | aDMan(1→2)aDMan(1→6)[aDMan(1→2)aDMan(1→3)]aDMan(1→6)<br>[aDMan(1→2)aDMan(1→2)aDMan(1→3)]bDMan(1→4)bDGlcNAc(1→4)bDGlcNAc(1→)PROA-339 |
| G13 | N355 | FA2 | 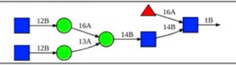 | bDGlcNAc(1→2)aDMan(1→6)[bDGlcNAc(1→2)aDMan(1→3)]bDMan(1→4)bDGlcNAc(1→4)<br>[aLFuc(1→6)]bDGlcNAc(1→)PROA-355 |
| G14 | N363 | M9 | 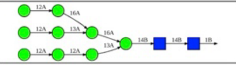 | aDMan(1→2)aDMan(1→6)[aDMan(1→2)aDMan(1→3)]aDMan(1→6)<br>[aDMan(1→2)aDMan(1→2)aDMan(1→3)]bDMan(1→4)bDGlcNAc(1→4)bDGlcNAc(1→)PROA-363 |
| G15 | N386 | M8 | 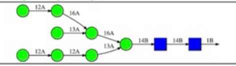 | aDMan(1→2)aDMan(1→6)[aDMan(1→3)]aDMan(1→6)<br>[aDMan(1→2)aDMan(1→2)aDMan(1→3)]bDMan(1→4)bDGlcNAc(1→4)bDGlcNAc(1→)PROA-386 |
| G16 | N392 | M9 | 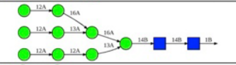 | aDMan(1→2)aDMan(1→6)[aDMan(1→2)aDMan(1→3)]aDMan(1→6)<br>[aDMan(1→2)aDMan(1→2)aDMan(1→3)]bDMan(1→4)bDGlcNAc(1→4)bDGlcNAc(1→)PROA-392 |
| G17 | N398 | M8 | 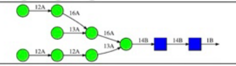 | aDMan(1→2)aDMan(1→6)[aDMan(1→3)]aDMan(1→6)<br>[aDMan(1→2)aDMan(1→2)aDMan(1→3)]bDMan(1→4)bDGlcNAc(1→4)bDGlcNAc(1→)PROA-398 |
| G18 | N406 | FA2G2S1 | 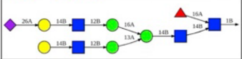 | aDNeu5Ac(2→6)bDGal(1→4)bDGlcNAc(1→2)aDMan(1→6)<br>[bDGal(1→4)bDGlcNAc(1→2)aDMan(1→3)]bDMan(1→4)bDGlcNAc(1→4)[aLFuc(1→6)]bDGlcNAc(1→)PROA-406 |
| G19 | N411 | M8 | 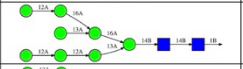 | aDMan(1→2)aDMan(1→6)[aDMan(1→3)]aDMan(1→6)<br>[aDMan(1→2)aDMan(1→2)aDMan(1→3)]bDMan(1→4)bDGlcNAc(1→4)bDGlcNAc(1→)PROA-411 |
| G20 | N416 | M9 | 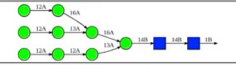 | aDMan(1→2)aDMan(1→6)[aDMan(1→2)aDMan(1→3)]aDMan(1→6)<br>[aDMan(1→2)aDMan(1→2)aDMan(1→3)]bDMan(1→4)bDGlcNAc(1→4)bDGlcNAc(1→)PROA-416 |
| G21 | N462 | FA2 | 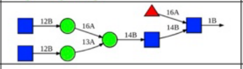 | bDGlcNAc(1→2)aDMan(1→6)[bDGlcNAc(1→2)aDMan(1→3)]bDMan(1→4)bDGlcNAc(1→4)<br>[aLFuc(1→6)]bDGlcNAc(1→)PROA-462 |
| G22 | N611 | A2G2 | 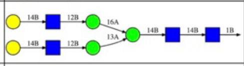 | bDGal(1→4)bDGlcNAc(1→2)aDMan(1→6)<br>[bDGal(1→4)bDGlcNAc(1→2)aDMan(1→3)]bDMan(1→4)bDGlcNAc(1→4)bDGlcNAc(1→)PROA-611 |
| G23 | N637 | FA2G2S1 | 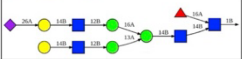 | aDNeu5Ac(2→6)bDGal(1→4)bDGlcNAc(1→2)aDMan(1→6)<br>[bDGal(1→4)bDGlcNAc(1→2)aDMan(1→3)]bDMan(1→4)bDGlcNAc(1→4)[aLFuc(1→6)]bDGlcNAc(1→)PROA-637 |

**Supplementary Table 2.** Glycan compositions for chain B. Glycosylation sites are reported using HXB2 notation.

CHAIN B

| # | SITE | TYPE | STRUCTURE | SEQUENCE |
|---|---|---|---|---|
| G1 | N88 | FA2G2S1 | 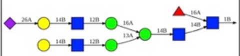 | aDNeu5Ac(2→6)bDGal(1→4)bDGlcNAc(1→2)aDMan(1→6)<br>[bDGal(1→4)bDGlcNAc(1→2)aDMan(1→3)]bDMan(1→4)bDGlcNAc(1→4)[aLFuc(1→6)]bDGlcNAc(1→)PROB-88 |
| G2 | N133 | FA2G2S1 | 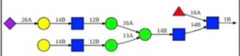 | aDNeu5Ac(2→6)bDGal(1→4)bDGlcNAc(1→2)aDMan(1→6)<br>[bDGal(1→4)bDGlcNAc(1→2)aDMan(1→3)]bDMan(1→4)bDGlcNAc(1→4)[aLFuc(1→6)]bDGlcNAc(1→)PROB-133 |
| G3 | N137 | FA2 | 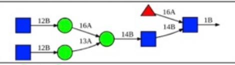 | bDGlcNAc(1→2)aDMan(1→6)[bDGlcNAc(1→2)aDMan(1→3)]bDMan(1→4)bDGlcNAc(1→4)<br>[aLFuc(1→6)]bDGlcNAc(1→)PROB-137 |
| G4 | N156 | M8 | 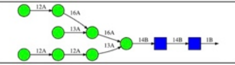 | aDMan(1→2)aDMan(1→6)[aDMan(1→3)]aDMan(1→6)<br>[aDMan(1→2)aDMan(1→2)aDMan(1→3)]bDMan(1→4)bDGlcNAc(1→4)bDGlcNAc(1→)PROB-156 |
| G5 | N160 | M8 | 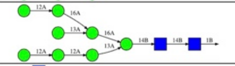 | aDMan(1→2)aDMan(1→6)[aDMan(1→3)]aDMan(1→6)<br>[aDMan(1→2)aDMan(1→2)aDMan(1→3)]bDMan(1→4)bDGlcNAc(1→4)bDGlcNAc(1→)PROB-160 |
| G6 | N197 | FA3 | 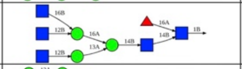 | bDGlcNAc(1→6)[bDGlcNAc(1→2)]aDMan(1→6)[bDGlcNAc(1→2)aDMan(1→3)]bDMan(1→4)bDGlcNAc(1→4)<br>[aLFuc(1→6)]bDGlcNAc(1→)PROB-197 |
| G7 | N234 | M9 | 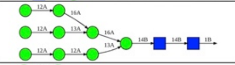 | aDMan(1→2)aDMan(1→6)[aDMan(1→2)aDMan(1→3)]aDMan(1→6)<br>[aDMan(1→2)aDMan(1→2)aDMan(1→3)]bDMan(1→4)bDGlcNAc(1→4)bDGlcNAc(1→)PROB-234 |
| G8 | N262 | M8 | 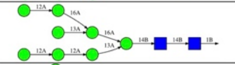 | aDMan(1→2)aDMan(1→6)[aDMan(1→3)]aDMan(1→6)<br>[aDMan(1→2)aDMan(1→2)aDMan(1→3)]bDMan(1→4)bDGlcNAc(1→4)bDGlcNAc(1→)PROB-262 |
| G9 | N276 | Hybrid G1 | 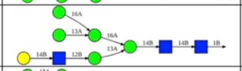 | bDGal(1→4)bDGlcNAc(1→2)aDMan(1→3)[aDMan(1→6)<br>[aDMan(1→3)]aDMan(1→6)]bDMan(1→4)bDGlcNAc(1→4)bDGlcNAc(1→)PROB-276 |
| G10 | N295 | M8 | 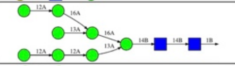 | aDMan(1→2)aDMan(1→6)[aDMan(1→3)]aDMan(1→6)<br>[aDMan(1→2)aDMan(1→2)aDMan(1→3)]bDMan(1→4)bDGlcNAc(1→4)bDGlcNAc(1→)PROB-295 |
| G11 | N301 | M9 | 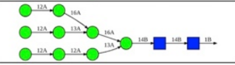 | aDMan(1→2)aDMan(1→6)[aDMan(1→2)aDMan(1→3)]aDMan(1→6)<br>[aDMan(1→2)aDMan(1→2)aDMan(1→3)]bDMan(1→4)bDGlcNAc(1→4)bDGlcNAc(1→)PROB-301 |
| G12 | N339 | M9 | 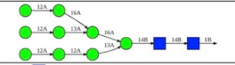 | aDMan(1→2)aDMan(1→6)[aDMan(1→2)aDMan(1→3)]aDMan(1→6)<br>[aDMan(1→2)aDMan(1→2)aDMan(1→3)]bDMan(1→4)bDGlcNAc(1→4)bDGlcNAc(1→)PROB-339 |
| G13 | N355 | FA3 | 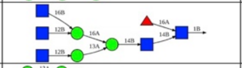 | bDGlcNAc(1→6)[bDGlcNAc(1→2)]aDMan(1→6)[bDGlcNAc(1→2)aDMan(1→3)]bDMan(1→4)bDGlcNAc(1→4)<br>[aLFuc(1→6)]bDGlcNAc(1→)PROB-355 |
| G14 | N363 | M8 | 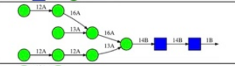 | aDMan(1→2)aDMan(1→6)[aDMan(1→3)]aDMan(1→6)<br>[aDMan(1→2)aDMan(1→2)aDMan(1→3)]bDMan(1→4)bDGlcNAc(1→4)bDGlcNAc(1→)PROB-363 |
| G15 | N386 | M8 | 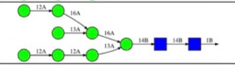 | aDMan(1→2)aDMan(1→6)[aDMan(1→3)]aDMan(1→6)<br>[aDMan(1→2)aDMan(1→2)aDMan(1→3)]bDMan(1→4)bDGlcNAc(1→4)bDGlcNAc(1→)PROB-386 |
| G16 | N392 | M9 | 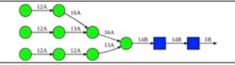 | aDMan(1→2)aDMan(1→6)[aDMan(1→2)aDMan(1→3)]aDMan(1→6)<br>[aDMan(1→2)aDMan(1→2)aDMan(1→3)]bDMan(1→4)bDGlcNAc(1→4)bDGlcNAc(1→)PROB-392 |
| G17 | N398 | M8 | 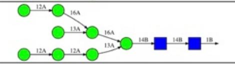 | aDMan(1→2)aDMan(1→6)[aDMan(1→3)]aDMan(1→6)<br>[aDMan(1→2)aDMan(1→2)aDMan(1→3)]bDMan(1→4)bDGlcNAc(1→4)bDGlcNAc(1→)PROB-398 |
| G18 | N406 | FA3G3 | 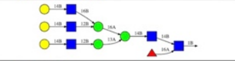 | bDGal(1→4)bDGlcNAc(1→6)[bDGal(1→4)bDGlcNAc(1→2)]aDMan(1→6)<br>[bDGal(1→4)bDGlcNAc(1→2)aDMan(1→3)]bDMan(1→4)bDGlcNAc(1→4)[aLFuc(1→6)]bDGlcNAc(1→)PROB-406 |
| G19 | N411 | M8 | 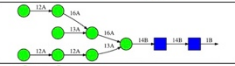 | aDMan(1→2)aDMan(1→6)[aDMan(1→3)]aDMan(1→6)<br>[aDMan(1→2)aDMan(1→2)aDMan(1→3)]bDMan(1→4)bDGlcNAc(1→4)bDGlcNAc(1→)PROB-411 |
| G20 | N416 | M9 | 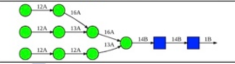 | aDMan(1→2)aDMan(1→6)[aDMan(1→2)aDMan(1→3)]aDMan(1→6)<br>[aDMan(1→2)aDMan(1→2)aDMan(1→3)]bDMan(1→4)bDGlcNAc(1→4)bDGlcNAc(1→)PROB-416 |
| G21 | N462 | FA3 | 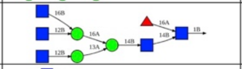 | bDGlcNAc(1→6)[bDGlcNAc(1→2)]aDMan(1→6)[bDGlcNAc(1→2)aDMan(1→3)]bDMan(1→4)bDGlcNAc(1→4)<br>[aLFuc(1→6)]bDGlcNAc(1→)PROB-462 |
| G22 | N611 | A3 | 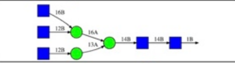 | bDGlcNAc(1→6)[bDGlcNAc(1→2)]aDMan(1→6)<br>[bDGlcNAc(1→2)aDMan(1→3)]bDMan(1→4)bDGlcNAc(1→4)bDGlcNAc(1→)PROB-611 |
| G23 | N637 | FA2 | 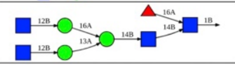 | bDGlcNAc(1→2)aDMan(1→6)[bDGlcNAc(1→2)aDMan(1→3)]bDMan(1→4)bDGlcNAc(1→4)<br>[aLFuc(1→6)]bDGlcNAc(1→)PROB-637 |

**Supplementary Table 3.** Glycan compositions for chain C. Glycosylation sites are reported using HXB2 notation.

CHAIN C

| # | SITE | TYPE | STRUCTURE | SEQUENCE |
|---|---|---|---|---|
| G1 | N88 | A2G2 | 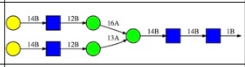 | bDGal(1→4)bDGlcNAc(1→2)aDMan(1→6) [bDGal(1→4)bDGlcNAc(1→2)aDMan(1→3)]bDMan(1→4)bDGlcNAc(1→4)bDGlcNAc(1→)PROC-88 |
| G2 | N133 | A2G2 | 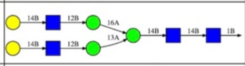 | bDGal(1→4)bDGlcNAc(1→2)aDMan(1→6) [bDGal(1→4)bDGlcNAc(1→2)aDMan(1→3)]bDMan(1→4)bDGlcNAc(1→4)bDGlcNAc(1→)PROC-133 |
| G3 | N137 | FA2 | 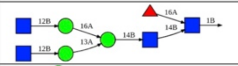 | bDGlcNAc(1→2)aDMan(1→6)[bDGlcNAc(1→2)aDMan(1→3)]bDMan(1→4)bDGlcNAc(1→4) [aLFuc(1→6)]bDGlcNAc(1→)PROC-137 |
| G4 | N156 | M7 | 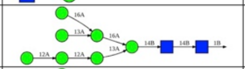 | aDMan(1→2)aDMan(1→2)aDMan(1→3)[aDMan(1→6) [aDMan(1→3)]aDMan(1→6)]bDMan(1→4)bDGlcNAc(1→4)bDGlcNAc(1→)PROC-156 |
| G5 | N160 | M7 | 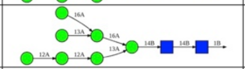 | aDMan(1→2)aDMan(1→2)aDMan(1→3)[aDMan(1→6) [aDMan(1→3)]aDMan(1→6)]bDMan(1→4)bDGlcNAc(1→4)bDGlcNAc(1→)PROC-160 |
| G6 | N197 | FA2G2S2 | 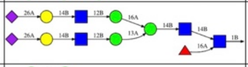 | aDNeu5Ac(2→6)bDGal(1→4)bDGlcNAc(1→2)aDMan(1→6) [aDNeu5Ac(2→6)bDGal(1→4)bDGlcNAc(1→2)aDMan(1→3)]bDMan(1→4)bDGlcNAc(1→4) [aLFuc(1→6)]bDGlcNAc(1→)PROC-197 |
| G7 | N234 | M9 | 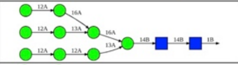 | aDMan(1→2)aDMan(1→6)[aDMan(1→2)aDMan(1→3)]aDMan(1→6) [aDMan(1→2)aDMan(1→2)aDMan(1→3)]bDMan(1→4)bDGlcNAc(1→4)bDGlcNAc(1→)PROC-234 |
| G8 | N262 | M8 | 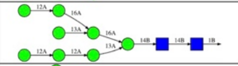 | aDMan(1→2)aDMan(1→6)[aDMan(1→3)]aDMan(1→6) [aDMan(1→2)aDMan(1→2)aDMan(1→3)]bDMan(1→4)bDGlcNAc(1→4)bDGlcNAc(1→)PROC-262 |
| G9 | N276 | Hybrid G1 | 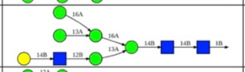 | bDGal(1→4)bDGlcNAc(1→2)aDMan(1→3)[aDMan(1→6) [aDMan(1→3)]aDMan(1→6)]bDMan(1→4)bDGlcNAc(1→4)bDGlcNAc(1→)PROC-276 |
| G10 | N295 | M8 | 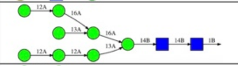 | aDMan(1→2)aDMan(1→6)[aDMan(1→3)]aDMan(1→6) [aDMan(1→2)aDMan(1→2)aDMan(1→3)]bDMan(1→4)bDGlcNAc(1→4)bDGlcNAc(1→)PROC-295 |
| G11 | N301 | M8 | 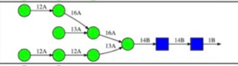 | aDMan(1→2)aDMan(1→6)[aDMan(1→3)]aDMan(1→6) [aDMan(1→2)aDMan(1→2)aDMan(1→3)]bDMan(1→4)bDGlcNAc(1→4)bDGlcNAc(1→)PROC-301 |
| G12 | N339 | M8 | 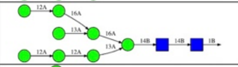 | aDMan(1→2)aDMan(1→6)[aDMan(1→3)]aDMan(1→6) [aDMan(1→2)aDMan(1→2)aDMan(1→3)]bDMan(1→4)bDGlcNAc(1→4)bDGlcNAc(1→)PROC-339 |
| G13 | N355 | Hybrid G1 | 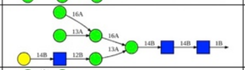 | bDGal(1→4)bDGlcNAc(1→2)aDMan(1→3)[aDMan(1→6) [aDMan(1→3)]aDMan(1→6)]bDMan(1→4)bDGlcNAc(1→4)bDGlcNAc(1→)PROC-355 |
| G14 | N363 | M8 | 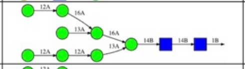 | aDMan(1→2)aDMan(1→6)[aDMan(1→3)]aDMan(1→6) [aDMan(1→2)aDMan(1→2)aDMan(1→3)]bDMan(1→4)bDGlcNAc(1→4)bDGlcNAc(1→)PROC-363 |
| G15 | N386 | M8 | 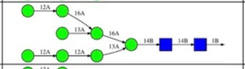 | aDMan(1→2)aDMan(1→6)[aDMan(1→3)]aDMan(1→6) [aDMan(1→2)aDMan(1→2)aDMan(1→3)]bDMan(1→4)bDGlcNAc(1→4)bDGlcNAc(1→)PROC-386 |
| G16 | N392 | M8 | 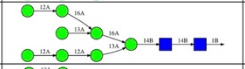 | aDMan(1→2)aDMan(1→6)[aDMan(1→3)]aDMan(1→6) [aDMan(1→2)aDMan(1→2)aDMan(1→3)]bDMan(1→4)bDGlcNAc(1→4)bDGlcNAc(1→)PROC-392 |
| G17 | N398 | M9 | 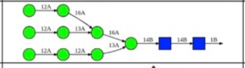 | aDMan(1→2)aDMan(1→6)[aDMan(1→2)aDMan(1→3)]aDMan(1→6) [aDMan(1→2)aDMan(1→2)aDMan(1→3)]bDMan(1→4)bDGlcNAc(1→4)bDGlcNAc(1→)PROC-398 |
| G18 | N406 | FA2 | 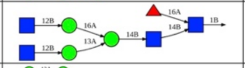 | bDGlcNAc(1→2)aDMan(1→6)[bDGlcNAc(1→2)aDMan(1→3)]bDMan(1→4)bDGlcNAc(1→4) [aLFuc(1→6)]bDGlcNAc(1→)PROC-406 |
| G19 | N411 | M9 | 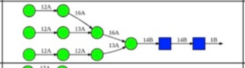 | aDMan(1→2)aDMan(1→6)[aDMan(1→2)aDMan(1→3)]aDMan(1→6) [aDMan(1→2)aDMan(1→2)aDMan(1→3)]bDMan(1→4)bDGlcNAc(1→4)bDGlcNAc(1→)PROC-411 |
| G20 | N416 | M8 | 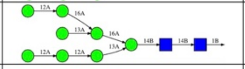 | aDMan(1→2)aDMan(1→6)[aDMan(1→3)]aDMan(1→6) [aDMan(1→2)aDMan(1→2)aDMan(1→3)]bDMan(1→4)bDGlcNAc(1→4)bDGlcNAc(1→)PROC-416 |
| G21 | N462 | FA2G2S2 | 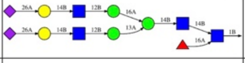 | aDNeu5Ac(2→6)bDGal(1→4)bDGlcNAc(1→2)aDMan(1→6) [aDNeu5Ac(2→6)bDGal(1→4)bDGlcNAc(1→2)aDMan(1→3)]bDMan(1→4)bDGlcNAc(1→4) [aLFuc(1→6)]bDGlcNAc(1→)PROC-462 |
| G22 | N611 | A2G2S2 | 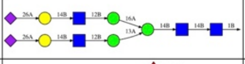 | aDNeu5Ac(2→6)bDGal(1→4)bDGlcNAc(1→2)aDMan(1→6) [aDNeu5Ac(2→6)bDGal(1→4)bDGlcNAc(1→2)aDMan(1→3)]bDMan(1→4)bDGlcNAc(1→4)bDGlcNAc(1→)PROC-611 |
| G23 | N637 | FA2 | 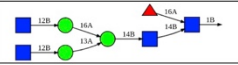 | bDGlcNAc(1→2)aDMan(1→6)[bDGlcNAc(1→2)aDMan(1→3)]bDMan(1→4)bDGlcNAc(1→4) [aLFuc(1→6)]bDGlcNAc(1→)PROC-637 |

**Supplementary Table 4.** Membrane lipid composition.

| Lipid Category | Lipid | Percent Abundance | Outer Leaflet Fraction | Outer Leaflet Count | Inner Leaflet Fraction | Inner Leaflet Count |
|---|---|---|---|---|---|---|
| PC | DPPC | 6.32 | 0.75 | 120 | 0.25 | 30 |
| | POPC | 9.48 | 0.75 | 150 | 0.25 | 46 |
| PE | POPE | 21.40 | 0.25 | 108 | 0.75 | 320 |
| PS | POPS | 13.72 | 0.00 | 0 | 1.00 | 260 |
| SM | PSM | 16.51 | 0.75 | 248 | 0.25 | 82 |
| Chl | CHOL | 32.56 | 0.60 | 416 | 0.40 | 260 |

**Supplementary Table 5.** Details of Simulated Systems.

| System | Box dimensions (Å x Å x Å) | Total # atoms | Salt Concentration (M) | R1 (ns) | R2 (ns) | R3 (ns) | R4 (ns) | R5 (ns) |
|---|---|---|---|---|---|---|---|---|
| Full length Env | 242 Å x 242 Å x 252 Å | 1297075 | 0.15 | 1050 | 1050 | 1050 | 1050 | 1050 |

**Supplementary Table 6.** Frequency of Glycan membrane contacts.

| Glycan/Chain | R1 | R2 | R3 | R4 | R5 | Average Frequency |
|---|---|---|---|---|---|---|
| **N88/A** | **0** | **0** | **0** | **60.46** | **87.32** | **29.56** |
| **N88/B** | **32.71** | **0.1** | **51.82** | **11.24** | **82.83** | **35.74** |
| **N88/C** | **25.07** | **0.08** | **84.66** | **0** | **0.02** | **21.96** |
| N133/A | 0 | 0 | 0 | 0 | 0 | 0 |
| N133/B | 0 | 0 | 0 | 0 | 0 | 0 |
| N133/C | 0 | 0 | 0 | 0 | 0 | 0 |
| N137/A | 0 | 0 | 0 | 0 | 0 | 0 |
| N137/B | 0 | 0 | 0 | 0 | 0 | 0 |
| N137/C | 0 | 0 | 0 | 0 | 0 | 0 |
| N156/A | 0 | 0 | 0 | 0 | 0 | 0 |
| N156/B | 0 | 0 | 0 | 0 | 0 | 0 |
| N156/C | 0 | 0 | 0 | 0 | 0 | 0 |
| N160/A | 0 | 0 | 0 | 0 | 0 | 0 |
| N160/B | 0 | 0 | 0 | 0 | 0 | 0 |
| N160/C | 0 | 0 | 0 | 0 | 0 | 0 |
| N197/A | 0 | 0 | 0 | 0 | 0 | 0 |
| N197/B | 0 | 0 | 0 | 0 | 0 | 0 |
| N197/C | 0 | 0 | 0 | 0 | 0 | 0 |
| N234/A | 0 | 0 | 0 | 3.39 | 50.05 | 10.69 |
| N234/B | 0.03 | 0 | 42.42 | 0 | 0 | 8.49 |
| N234/C | 0 | 0 | 0.02 | 0 | 0 | 0.004 |
| N262/A | 0 | 0 | 0 | 0 | 0 | 0 |
| N262/B | 0 | 0 | 0 | 0 | 0 | 0 |
| N262/C | 0 | 0 | 0 | 0 | 0 | 0 |
| N276/A | 0 | 0 | 0 | 1.3 | 3.37 | 0.93 |
| N276/B | 0 | 0 | 16.31 | 0 | 0 | 3.26 |
| N276/C | 0 | 0 | 0 | 0 | 0 | 0 |
| N295/A | 0 | 0 | 0 | 14.94 | 0.51 | 3.09 |
| N295/B | 0 | 0 | 0 | 0 | 0.21 | 0.04 |
| N295/C | 0 | 0 | 30.57 | 0 | 0 | 6.11 |
| N301/A | 0 | 0 | 0 | 0 | 0 | 0 |
| N301/B | 0 | 0 | 0 | 0 | 0 | 0 |
| N301/C | 0 | 0 | 0 | 0 | 0 | 0 |
| N339/A | 0 | 0 | 0 | 0.06 | 0 | 0.01 |
| N339/B | 0 | 0 | 0 | 0 | 0 | 0 |
| N339/C | 0 | 0 | 0 | 0 | 0 | 0 |

| | | | | | | |
|---|---|---|---|---|---|---|
| N355/A | 0 | 0 | 0 | 33.38 | 19.91 | 10.66 |
| N355/B | 0 | 0 | 0 | 0 | 0 | 0 |
| N355/C | 0 | 0 | 28.55 | 0 | 0 | 5.71 |
| N363/A | 0 | 0 | 0 | 0 | 0 | 0 |
| N363/B | 0 | 0 | 0 | 0 | 0 | 0 |
| N363/C | 0 | 0 | 0 | 0 | 0 | 0 |
| N386/A | 0 | 0 | 0 | 0 | 0 | 0 |
| N386/B | 0 | 0 | 0 | 0 | 0 | 0 |
| N386/C | 0 | 0 | 0 | 0 | 0 | 0 |
| N392/A | 0 | 0 | 0 | 0 | 0 | 0 |
| N392/B | 0 | 0 | 0 | 0 | 0 | 0 |
| N392/C | 0 | 0 | 0 | 0 | 0 | 0 |
| N398/A | 0 | 0 | 0 | 29.42 | 0.84 | 6.05 |
| N398/B | 0 | 0 | 1.97 | 0 | 0 | 0.39 |
| N398/C | 0 | 0 | 10.27 | 0 | 0 | 2.05 |
| N406/A | 0 | 0 | 0 | 0 | 0 | 0 |
| N406/B | 0 | 0 | 0 | 0 | 0 | 0 |
| N406/C | 0 | 0 | 25.54 | 0 | 0 | 5.11 |
| N411/A | 0 | 0 | 0 | 2.5 | 0 | 0.5 |
| N411/B | 0 | 0 | 0 | 0 | 0 | 0 |
| N411/C | 0 | 0 | 26.21 | 0 | 0 | 5.24 |
| N416/A | 0 | 0 | 0 | 0.01 | 0 | 0 |
| N416/B | 0 | 0 | 0 | 0 | 0 | 0 |
| N416/C | 0 | 0 | 12.58 | 0 | 0 | 2.52 |
| N462/A | 0 | 0 | 0 | 1.1 | 1.54 | 0.53 |
| N462/B | 0 | 0 | 0 | 0 | 0 | 0 |
| N462/C | 0 | 0 | 0.7 | 0 | 0 | 0.14 |
| **N611/A** | **2.19** | **0.96** | **0.13** | **73.3** | **92.73** | **33.86** |
| **N611/B** | **66.24** | **0.63** | **89.43** | **0** | **0.01** | **31.26** |
| **N611/C** | **0.48** | **0.56** | **1.3** | **48.39** | **62.72** | **22.69** |
| N637/A | 0 | 0 | 0 | 39.23 | 9.99 | 9.84 |
| N637/B | 0.17 | 0 | 41.71 | 0 | 0 | 8.38 |
| N637/C | 0 | 0 | 0 | 0 | 0 | 0 |

**Supplementary Table 7.** HIV-1 Env's major immunogenic regions. Residue masks are provided using the UniProt numbering scheme, which is consistent with the residue numbering of the Env model simulated in this study. For conversion to HXB2 notation, please refer to the alignment in Figure S1. Glycans are reported using HXB2 notation.

| Epitope | Antibody | Residue masks | Reference |
|---|---|---|---|
| **gp120-gp41 interface** | PGT151 | 46–56, 480–494, 503–511<br>glycan N611 | 52 |
| | 8ANC195 | 14–19, 59–64, 207–209, 580–584, 596–606 | 53 |
| | 35O22 | 59–64, 207–209, 586–593, 596–603<br>glycan N88 | 54 |
| **V1V2** | PGT145 | 94–96, 121–124, 127–131,<br>3 x glycan N160 | 55 |
| | PGDM1400 | 94–96, 121–124, 127–131,<br>3 x glycan N160 | 56 |
| | PG9 | 127–135,<br>glycan N156, glycan N160 | 57 |
| | PG16 | 127–135, 151–157,<br>glycan N156, glycan N160 | 58 |
| **V3** | PGT121 (V3base glycan supersite +V1) | 292–299 and V1's 104–107,<br>glycan N332, glycan N137 | 55 |
| | BG18 (V3base glycan supersite +V1) | 292–299,<br>glycan N332, glycan N137 | 59 |
| | M4008_1 (V3crown) | 271–285, 291–293 | 60 |
| **CD4 binding site** | VRC01 | 333–341, 397–400, 423–430, 435–44,<br>glycan N276 | 61 |
| | CH103 | 333–341, 397–400, 423–430, 435–443,<br>glycan N276 | 62 |
| | N6 | 333–341, 397–400, 423–430, 435–443,<br>glycan N276 | 63 |
| **Fusion Peptide** | N123-VRC34.01 | 480–494 (HXB2: 512–526) | 64 |
| | PGT151 | 480–494 (HXB2: 512–526) | 52 |
| **Silent Face** | VRC-PG05 | 260–263, 303–306, 413–416,<br>glycan N262, glycan N295, glycan N448 | 65 |
| | SF12 | 27–29, 183–184, 222, 260–263, 413–416,<br>glycan N262, glycan N295, glycan N448 | 66 |
| **MPER** | 10E8 | 639–651 (HXB2: 671–683) | 14 |
| | 4E10 | 639–651 (HXB2: 671–683) | 67 |
| | Z13e1 | 634–645 (HXB2: 666–677) | 68 |
| | 2F5 | 624–639 (HXB2: 656–671) | 69 |

### 4. Captions for Supplementary Movies

**Supplementary Movie 1.** All-atom MD simulation of the full-length glycosylated HIV-1 Env trimer embedded in a biologically relevant lipid bilayer. ~1 μs of simulation is shown, illustrating significant tilting of the Env trimer relative to the membrane normal. N-linked glycans are depicted as dynamic blue meshes, the protein backbone is shown in light blue, and the viral membrane is represented with purple van der Waals (vdW) spheres.
